# Antigen-scaffolds drive preferential expansion of functional genetically engineered CAR and TCR T cells

**DOI:** 10.1101/2025.11.01.686037

**Authors:** Kristoffer Haurum Johansen, Keerthana Ramanathan, Carlos Rodriguez-Pardo, Anne Kargaard Gjelstrup, Hólmfridur Rósa Halldórsdóttir, Lasse Frank Voss, Siri Tvingsholm, Laura Stentoft Grand, Edoardo Dionisio, Alberte H. Skriver, Louise Pii Frederiksen, Mette Bjørnson, Marcus Svensson-Frej, Maria Ormhøj, Sine Reker Hadrup

## Abstract

The engineering of autologous T cells to express chimeric antigen receptors (CARs) can induce profound clinical responses in haematological malignancies, while T cell receptor-engineered T (TCR T) cells have led to durable responses in clinical trials for solid tumours. However, clinical manufacturing of engineered T cells is resource-intensive and often yields highly differentiated, exhausted effector T cells. To circumvent this, we have developed an antigen-scaffold (Ag-scaffold) technology to preferentially expand genetically engineered T cells. Such Ag-scaffolds present cognate antigen together with stimulatory factors such as cytokines. By providing a specific and receptor-engaging stimulation to CAR/TCR T cells, the expanded product is highly enriched for engineered T cells with a favourable proliferative and efficacious phenotype.

We expanded TCR T cells with Ag-scaffolds presenting peptide MHC (pMHC), and anti-CD19 CAR T cells with Ag-scaffolds presenting CD19 antigen. By applying cognate pMHC Ag-scaffolds, we achieved >80% antigen-specific T cells (83.62%±9.2%) after 14 days of culture with a distinct cytotoxic, proliferative phenotypical profile. Ag-scaffold expansion enhanced initial TCR and CAR cytotoxicity; sustained control was observed after repeated rechallenges of CAR T cells. *In vivo*, Ag-scaffold-expanded CRISPR/Cas9-engineered anti-CD19 CAR T also showed complete tumour eradication in a B-cell lymphoma xenograft model with a low dose of CAR T cells, which was not achieved using IL2/7/15 expansion.

## Introduction

T-cell engineering aimed at modifying the T-cell specificity using synthetic antigen receptors, such as chimeric antigen receptors (CARs) or recombinant T-cell receptors (TCRs), has demonstrated significant efficacy in treating malignancies^1^. CAR-engineered T cells (CAR T cells) can induce durable responses in hematologic malignancies; however, they have been less successful in treating solid cancers^2^. With the approval of the first FDA TCR-engineered T-cell therapy, afamitresgene autoleucel (Tecelra®), targeting MAGE-A4/HLA-A*02:01^+^ synovial sarcomas, TCR therapies have shown efficacy in treating solid malignancies^2^.

Traditionally, synthetic antigen receptors are introduced into T cells *ex vivo* by lentiviral transduction, followed by expansion with a cytokine cocktail of IL2, IL7, and IL15^1,3^. However, in recent years, pre-clinical and clinical studies have investigated nonviral introduction of CARs and TCRs into the endogenous TCR alpha constant locus (TRAC) by CRISPR/Cas9-mediated knockin (KI), allowing for endogenous control of the synthetic antigen receptor expression whilst simultaneously replacing the endogenous TCR^4,5^. Pre-clinical evidence supports that expressing the CAR/TCR under the regulation of the endogenous TCR promoter enhances the efficacy of the anti-tumour effect of the T cells^5,6^.

Although engineered T-cell therapies have proven successful in cancer immunotherapy, their efficacy is governed by the phenotype and functional properties of the final adoptive cell therapy product (ACT). Long-term follow-up of the CD19 CAR T cell-treated patients shows that, despite the initial complete response, a significant proportion of patients experience relapse due to poor cytotoxicity and persistence of the CAR T cells^7–9^. The antigen stimulus and the cytokines present during the *ex vivo* expansion greatly influence T-cell phenotype and functional properties. Early differentiated T-cell populations, such as stem-cell memory (T_SCM_) and central memory (T_CM_), have demonstrated enhanced tumour control in pre-clinical studies and early clinical trials due to their high self-renewal capacity^7,10,11^. On the contrary, the more differentiated T-cell phenotypes, such as T_EMRA_ and T_reg_ cells, are associated with reduced therapeutic efficacy due to their poor persistence and functionality^12^. The conventional *ex vivo* expansion protocol involves initial stimulation with anti-CD3/anti-CD28 antibodies, followed by expansion in IL2-supplemented media^13,14^. However, the unspecific polyclonal activation driven by anti-CD3/CD28 and IL2 leads to a highly differentiated T-cell phenotype, resulting in a suboptimal ACT product^15^. Consequently, subsequent studies have identified that the regulation of early differentiated phenotypes depends on other cytokines that share the common gamma chain, namely IL7, IL15, and IL21^3,16, 17^.

Additionally, antigen-mediated activation and co-stimulation using autologous (monocyte-derived dendritic cells; moDC) or artificial antigen-presenting platforms in combination with cytokines have shown some pre-clinical promise in improving the ACT product^14,18^. While antigen-specific T-cell expansion using patient-derived dendritic cells (DCs) has shown clinical efficacy, challenges remain in the complex manufacturing process. The recent advances in generating off-the-shelf, customisable, and biocompatible nano- and micro-materials have facilitated the development of novel antigen-presenting platforms that can provide T cells with the optimal signals to support the phenotype and functional characteristics of a desirable ACT product^18,19^.

Here, we apply a linear streptavidin-coated dextran scaffold that can be customised by coupling the respective biotinylated CAR antigen or pMHC complex, together with cytokines (Ag-scaffold), to specifically activate and expand TCR or CAR T cells for ACT^20^. We applied IL-2/IL-21 on the Ag-scaffold based on earlier optimisation showing superiority of this cytokine combination in antigen-specific T-cell expansion^20^ and the concept that Ag-scaffold-mediated antigen engagement fosters localised paracrine cytokine signalling. We tested the Ag-scaffold technology in T cells engineered with CRISPR/Cas9 KI or lentiviral vectors and compared their expansion and phenotype extensively with conventional expansion protocols. We demonstrate that Ag-scaffold-mediated expansion of engineered T cells yields an enriched T-cell product characterised by high self-renewal, enhanced proliferation, and increased cytotoxicity.

## Results

### Ag-scaffolds specifically enrich for genetically engineered TCR and CAR T cells

Non-viral CRISPR/Cas9-mediated KI of TCRs and CARs in the TRAC locus is relatively low (10-30%) (**Figure S1A-C**) (gRNA and HDRT sequences in Table S1). To test whether we could specifically expand T cells with successful KI, we established a method using Ag-scaffolds presenting the pMHC antigen for TCRs (TCR Ag-scaffolds) or CAR antigen for CARs (CAR Ag-scaffolds) together with IL2 and IL21, as shown to be the optimal Ag-scaffold composition in earlier work^21^. We knocked in TCRs or CARs 2 days after T-cell activation and expanded the TCR or CAR T cells with Ag-scaffolds added to the culture on days 4, 7, and 10, followed by analysis on day 14 (**Figures 1A-B**). To test whether we could use TCR Ag-scaffolds to specifically expand genetically engineered TCR T cells, we knocked in the alpha and beta chains of the NY-ESO-1-specific (SLLMWITQC/HLA-A*02:01) 1G4 TCR in the TRAC locus and compared expansion of T cells with soluble IL2/IL7/IL15 to Ag-scaffold expansion using SLLMWITQC/HLA-A*02:01/IL2/IL21-coated dextran scaffolds (TCR Ag-scaffolds). Using TCR Ag-scaffolds, CRISPR/Cas9 NY-ESO-1-specific TCR T cells were highly enriched (83.62%±9.2%) compared to T cells only expanded with cytokines (18.5%±9.2%) (**Figures 1C-E**). Further, the total number of antigen-specific TCR T cells was increased in the TCR Ag-scaffold-expanded population, with a 160-fold (SD=35.08) expansion of the TCR T cells after Ag-scaffold feeding, compared to 39-fold (SD=4.48) expansion in the cytokine controls over the same timeframe (**Figures 1F-H**). Interestingly, we did not observe specific expansion until after the second feed on day 5 post-KI **(Figures 1D, H**), suggesting that initial activation drove proliferation in the first days of culture.

**Figure 1.**
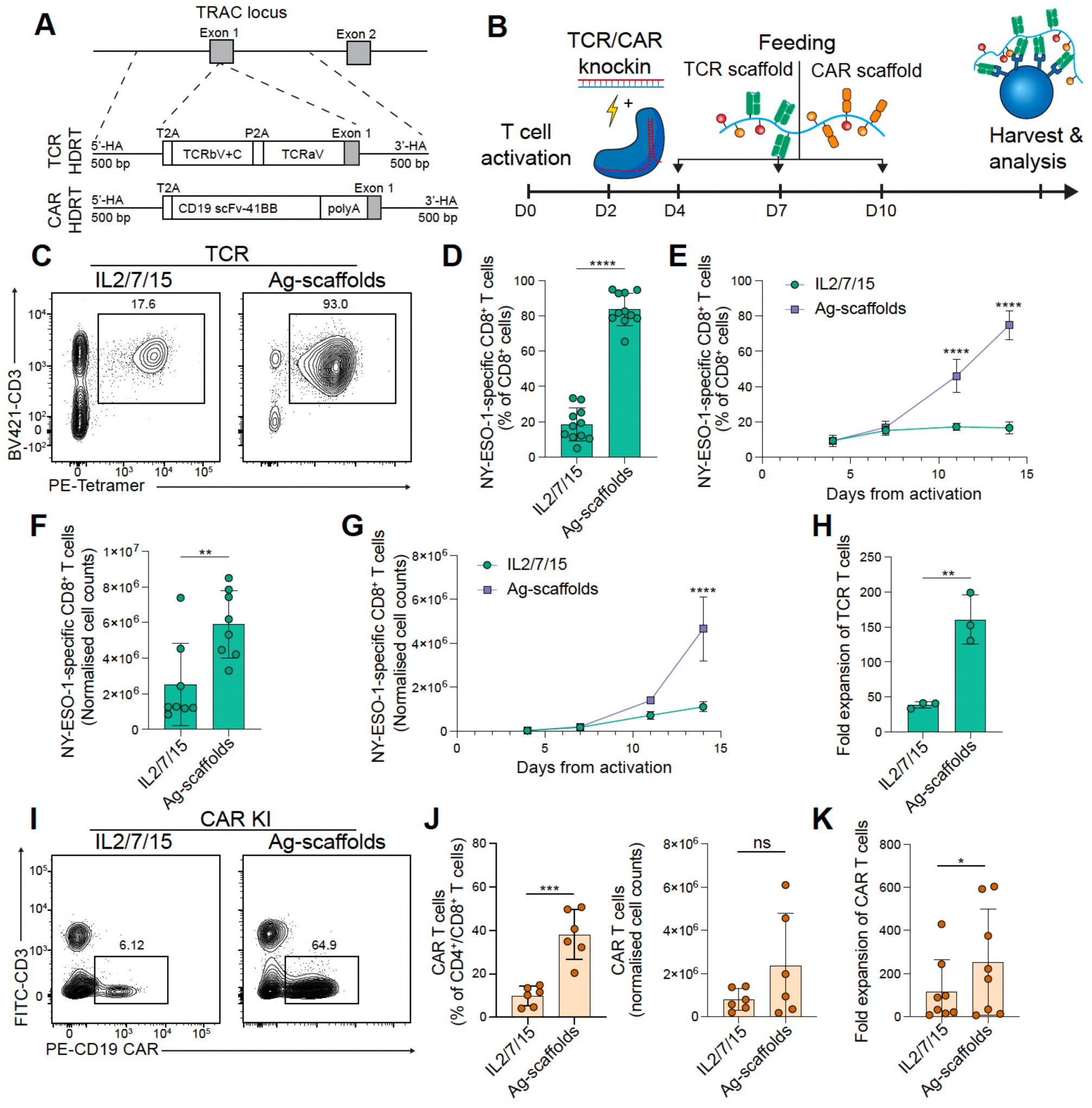
Antigen scaffold-mediated expansion of TCR- and CAR T cells greatly enriches KI TCR and CAR T cells. (**A**) Schematic representation of TCR and CAR KI homology-directed repair template (HDRT) with 500 bp homology arms (HA) targeting exon 1 of the TRAC locus. (**B**) A timeline illustrating the activation, KI, and Ag-scaffold feeding of engineered T cells. (**C**) Representative flow cytometry plots of NY-ESO-1-specific CRISPR/Cas9 1G4 TCR T cells after expansion with either IL2/IL7/IL15 or Ag-scaffolds. (**D-E**) Comparison of tetramer-positive (Tet^+^) percentages of CD8^+^ T cells at day 14 (**D**; n = 11 biological replicates per group), and (**E**) at different timepoints during expansion (n = 3 biological replicates per group). (**F-H**) Comparison of number of total specific cells at day 14 (**F**; n = 8 biological replicates per group), and over time (**G**) as well as the fold expansion from day 4 normalised to an initial count of 1 million total cells (**G, H**; n = 3 biological replicates per group). (**I**) Representative flow cytometry plots of CAR T cells expanded with IL2/7/15 or Ag-scaffolds. (**J**) Comparison of tetramer-positive (Tet^+^) percentages of CD4^+^/CD8^+^ T cells (n = 6 biological replicates per group) and cell counts (n = 6 biological replicates per group) following Ag-scaffold or IL2/IL7/IL15 expansion, with (**K**) fold expansion calculated from day 4 normalised to an initial count of 1×10^6^ total cells. Data in panels (**D-F**) represent pooled results from 3-4 experiments, (**H**) and (**K**) represent pooled results from 2 experiments presented as mean ± SD, with each dot indicating one PBMC sample post-KI. Statistical significance determined by ratio-paired t-test: *P < 0.05, **P < 0.01, ***P < 0.001, ****P < 0.0001.

Similarly to TCR KI, CAR KI in the TRAC locus exhibits limited KI efficiency (**Figure S1C**). We therefore generated CAR Ag-scaffolds using biotinylated CD19 antigen together with IL2 and IL21 to expand anti-CD19 CAR T cells. The CD19 CAR was knocked into T cells using the protocol described above. The KI CAR T cells were fed with CAR Ag-scaffolds every 3-4 days from day 4 post-activation as described above. The scaffolds were tested with 10 µM, 25 µM, and 50 µM per million T cells, and the results demonstrate a specific expansion effect using the CAR Ag-scaffold, which saturates at 25 µM/10^6^ cells (**Figure S1D**). Hence, we chose to continue with 25 uM of Ag-scaffolds per million T cells for CAR Ag-scaffolds. CAR Ag-scaffold expansion of CRISPR/Cas9 KI CAR T cells yielded a product with >35% CAR T cells (38.1%±11.5%) (**Figure 1I-J).** Furthermore, the CAR Ag-scaffold expansion resulted in a superior 400-fold expansion of CAR T cells, compared to 190-fold using free soluble cytokines (**Figure 1K**).

### Ag-scaffolds drive a distinct transcriptomic T cell phenotype with high proliferative capacity

To evaluate the effects of Ag-scaffold expansion on the phenotype and functional capacity of lentiviral or CRISPR/Cas9-engineered CAR T or TCR T cells, we compared the use of Ag-scaffolds to several established expansion protocols. Specifically, we tested two conventional cytokine cocktails commonly used for expanding engineered T cells: IL2 (200 IU/mL) + IL7 (5 ng/mL) + IL15 (5 ng/mL), and IL7 (5 ng/mL) + IL15 (5 ng/mL). Additionally, we evaluated IL2 (20 IU/mL) + IL21 (30 ng/mL) in solution to directly compare the effects of soluble versus scaffold-bound cytokines **(Figures S2A-D).** We proceeded with in-depth comparisons of the scaffold-expanded T cells to the cytokine cocktail (IL2 (200 IU/ml) + IL7 (5 ng/mL) + IL15 (5 ng/mL)) for further phenotypic and extensive functional analysis.

The expansion of engineered CAR T or TCR T cells yielded a product predominantly consisting of terminally differentiated T_EMRA_ cells post-expansion in both cytokine and Ag-scaffold conditions **(Figure 2A, Figures S2E-H)**. The expanded CAR T and TCR T cells had significantly higher percentages of CD27⁺CD28⁺ cells and lower percentages of CD27^-^CD28^-^cells, indicative of enhanced self-renewal capacity **(Figures 2B-C, Figures S2I-J**)^22^. Notably, the TCR Ag-scaffold-expanded T cells also had a significantly higher percentage of PD1^+^LAG3^+^ T cells compared to IL2/7/15-expanded TCR T cells. This difference was not observed in CAR Ag-scaffold-expanded CAR T cells **(Figure 2D, Figures S2K-I**).

**Figure 2.**
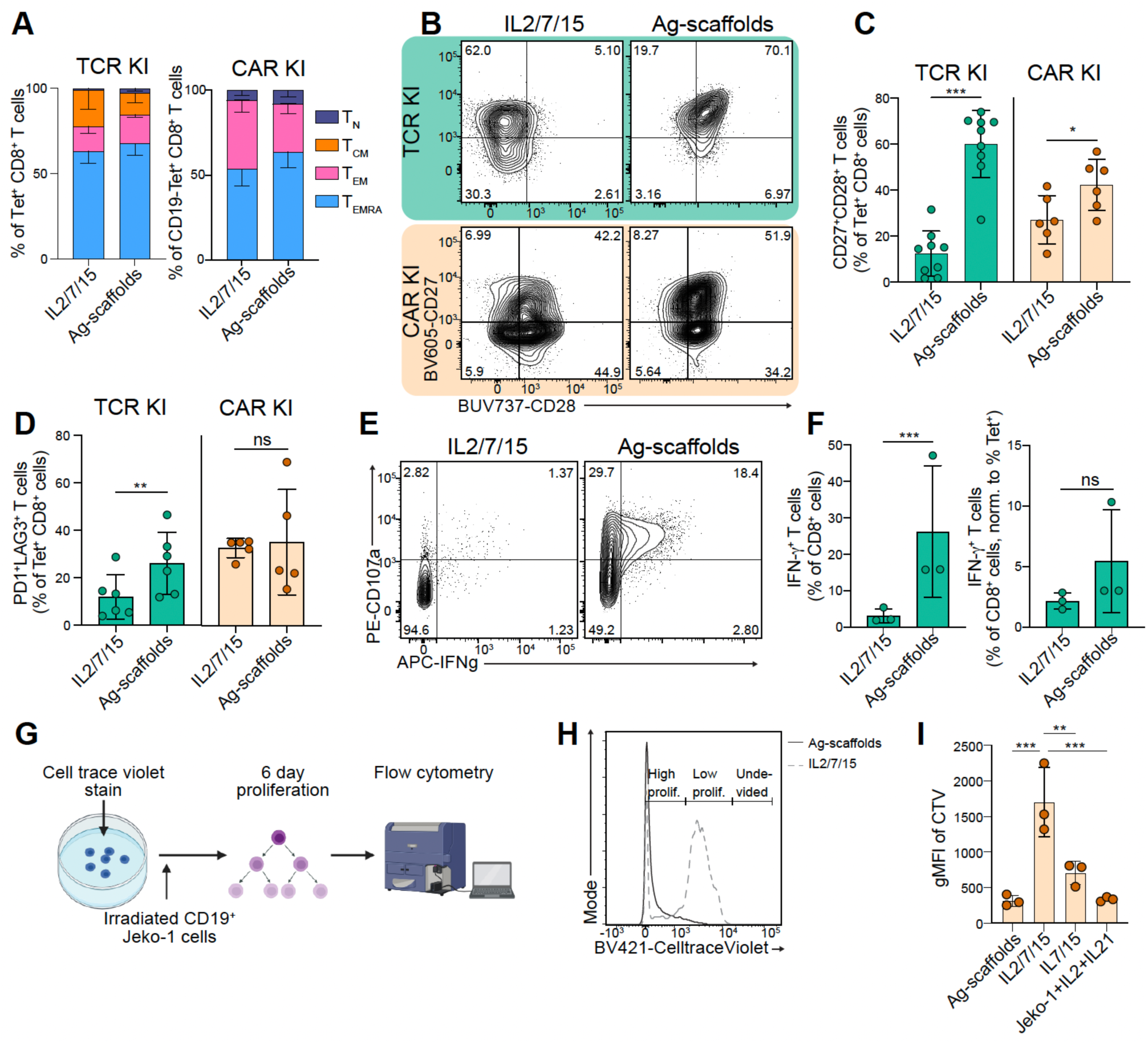
Ag-scaffold-mediated expansion of CRISPR/Cas9 T cells yields phenotypically distinct cells with superior cytotoxicity. (**A-F**) Phenotypic and functional analysis of CRISPR/Cas9 TCR/CAR KI T cells expanded using CAR/TCR Ag-scaffolds or cytokine cocktail (IL2/7/15). (**A**) Phenotypic distribution of KI TCR/CAR T cells based on CCR7 and CD45RA expression, showing T_N_ (CD45RA^+^CCR7^+^), T_CM_ (CD45RA^-^CCR7^+^), T_EM_, and T_EMRA_ subsets. (**B)** CRISPR/Cas9 KI CD8^+^Tet^+^ T cells expressing CD27^+^CD28^+^ with BV605-CD27 on y-axis and BUV737-CD28 on x-axis. **(C)** Graph showing the % CD27^+^CD28^+^ of CD8^+^Tet^+^ TCR/CAR KI T cells. (**D**) Graph showing the % of CD8^+^ Tet^+^ T cells expressing PD1^+^LAG3^+^ double positives. **(E-F)** Graph showing the expression % of TCR T cells expressing CD107a and IFN-γ when re-stimulated with target cells for 6 hrs. **(E)** Representative plot showing CRISPR/Cas9 KI CD8^+^Tet^+^ TCR T cells expressing CD107a^+^ IFN-γ^+^ with PE-CD107a on y-axis and APC-IFN-γ on x-axis. **(F)** Graph showing the expression % of T cells expressing IFN-γ, when re-stimulated with target cells for 6 hrs (left plot) normalised to the number of NY-ESO-1-specific 1G4 TCR^+^ T cells respectively (right plot). **(G)** Proliferative capacity of expanded CRISPR/Cas9 KI CD19 CAR T cells, as measured by loss of Cell Trace Violet in a 6-day co-culture with irradiated Jeko-1 target cells at an Effector: Target ratio of 1:5. **(H)** Representative histogram of cell trace violet expression of CAR+ T cells on day 6 of restimulation. **(I)** Graph showing geometric MFI of the cell trace violet on day 6. Data in (**B, F, H, I**) is from one experiment, and data in (**C**, **D**) is from 2-3 experiments presented as mean ± SD and statistical significance determined by ratio-paired t-tests: *P < 0.05, **P < 0.01, ***P < 0.001, ****P < 0.0001.

To investigate the functional capacity of the expanded cell product, we incubated expanded 1G4 TCR T cells with NY-ESO-1^+^/HLA-A*02:01^+^ A375 melanoma cells and evaluated production of IFN-γ, TNF-α, and presentation of the CD107a marker of degranulation by intracellular cytokine staining. A larger percentage of Ag-scaffold-expanded cells were positive for both IFN-γ, CD107a, and TNF-α (**Figure 2F**, **Figures S3A-F**). This is likely a consequence of the larger proportion of the T cells having the engineered TCR, as normalisation to the fraction of antigen-specific cells reduced this difference (**Figures 2E-F, Figures S3G-I**). Together, these results indicated that the Ag-scaffold-expanded T cells were at least on par with cytokine-expanded T cells in terms of producing IFN-γ and TNF-α.

Given the high levels of CD28 and CD27 following Ag-scaffold expansion, we evaluated the proliferative potential of Ag-scaffold-expanded CAR T cells and cytokine-expanded CAR T cells by co-incubating CellTrace Violet-stained CAR T cells with CD19^+^ Jeko-1 cells for 6 days followed by evaluation of CellTrace Violet dilution (**Figure 2G**). We found that Ag-scaffold- and IL2/21-expanded CAR T cells proliferated the most following *in vitro* Jeko-1 challenge in accordance with the high CD28/CD27 expression (**Figures 2H-I, Figures S3M-N**). Together, this indicates that Ag-scaffold-expanded cells are effective at producing cytokines and possess a high proliferative capacity.

Given the phenotypical differences and increased proliferative capacity observed following Ag-scaffold expansion, we next evaluated the transcriptional differences between sorted Ag-scaffold-and cytokine-expanded NY-ESO-1-specific TCR T cells using single-cell RNA sequencing **(Figure 3A)** and bulk RNA sequencing **(Figures S4A-B)**. Unsupervised clustering of single-cell transcriptomes revealed distinct separation of CRISPR-engineered Ag-scaffold-expanded TCR T cells from their cytokine-expanded counterparts **(Figure 3B)** with similar findings for lentiviral-engineered TCR T cells **(Figure S5A)**, indicating broad transcriptional reprogramming driven by the expansion method. We observed the largest differences in gene expression when comparing Ag-scaffold-expanded cells and IL2/7/15 cells, with the negative regulators of TCR signalling, SOCS1 and CISH^23,24^, being depleted in Ag-scaffold-expanded T cells **(Figures 3C-D, Figures S5B-C)**. We further observed an elevated expression of proliferation-associated genes, including PCNA, IL2RA, and MKI67, in Ag-scaffold-expanded T cells compared to both cytokine-expanded conditions, consistent with the high proliferative potential of Ag-scaffold-expanded T cells **(Figures 3C-E)**. In line with this observation, cell-cycle analysis demonstrated that approximately 50% of Ag-scaffold-expanded TCR T cells were in S or G2/M phase after 14 days of culture, confirming sustained cell-cycle activity **(Figure 3F, Figures S5D**).

**Figure 3.**
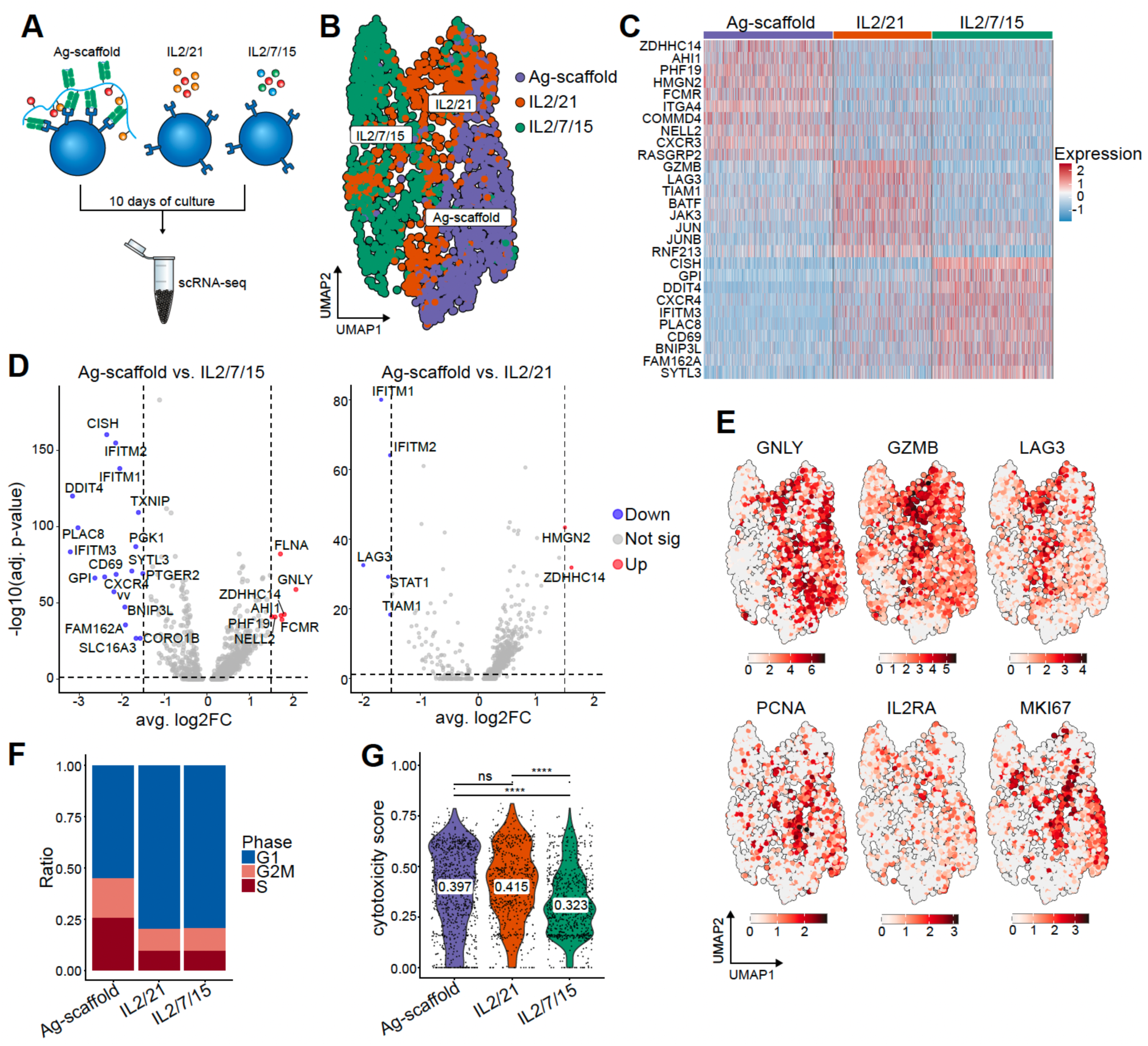
Single-cell RNA sequencing (scRNAseq) of CRISPR/Cas9-engineered T cells expanded with either Ag-scaffold, IL2/7, or IL2/7/15. (**A**) Diagram illustrating TCR T cells expanded with Ag-scaffold, IL2/21, or IL2/7/15 followed by 10x scRNAseq. (**B**) Uniform Manifold Approximation and Projection (UMAP) plot of all CRISPR/Cas9 engineered TCR T cells (total; 2,292 cells), with clusters colored and annotated according to cells expanded with either Ag-scaffold (total; 850 cells), IL2/21 (total; 647 cells), or IL2/7/15 (total; 795 cells). (**C**) Volcano plots showing differentially expressed genes for cells expanded with Ag-scaffold compared to IL2/7/15 (left) or IL2/21 (right). Genes significantly up- or down-regulated in the Ag-scaffold-expanded cluster are highlighted in red and blue, respectively. Dotted lines indicate thresholds on significance (log-fold-change =< 1.5 and p-value < 0.05 (-log10(0.05) = 1.3)). (**D**) UMAP plot of all CRISPR/Cas9 engineered TCR T cells showing the single cell mRNA expression levels of GNLY, GZMB, LAG3, PCNA, IL2RA, MKI67. (**E**) Expression of selected genes plotted on the UMAP from (D). (**F**) Barplot showing the ratio of cells in each cell cycle phase (G1, S, G2/M) across the three expansion conditions. (**G**) Violin plot illustrating cytotoxicity gene signature scores and means across the expansion conditions, calculated from RNA expression of selected markers using UCell (GZMA, GZMB, GNLY, PRF1, ITGB1, LAMP1). Statistics are calculated using Wilcoxon signed-rank test: ns = not significant, *P < 0.05, **P < 0.01, ***P < 0.001, ****P < 0.0001.

Differential gene expression analysis further revealed upregulation of AHI1 and HMGN2, which are implicated in cytokine-mediated proliferative signalling. In addition, Ag-scaffold-expanded T cells displayed marked increases in cytotoxic effector transcripts such as GZMB, GNLY, and PRF1 compared to IL2/7/15-expanded T cells **(Figures 3C-E, Figures S5E-G)**. This pattern was also observed in IL2/21-expanded cultures, suggesting that IL-2/21 signalling contributes to this cytotoxic gene program **(Figure 3G)**. We observed higher levels of LAG3 in IL2/21-expanded T cells compared to Ag-scaffold- and IL2/7/15-expanded T cells, suggesting a slightly more terminal phenotype in these cells **(Figure 3E)**. Together, these findings indicate that Ag-scaffold expansion drives a proliferative and cytotoxic effector phenotype distinct from that induced by soluble cytokine expansion. Similar phenotypic patterns were observed following bulk RNA sequencing of Ag-scaffold-expanded TCR T cells **(Figures S4A-B)**.

### Pooled tumour-shared antigen-specific CRISPR TCR T cells are effectively enriched by Ag-scaffolds

For TCR-based therapeutics, a challenge is the risk of tumour antigen-escape. To mitigate this, a pool of TCRs could be used to target several cancer antigens. We therefore tested the ability of the Ag-scaffold technology to stimulate and expand a pool of engineered TCR specificities. We engineered T cells by CRISPR/Cas9 KI of 5 HLA-A*02:01-restricted tumour-shared antigen-specific TCRs from the literature targeting NY-ESO-1/SLLMWITQC, MART-1/EAAGIGLTV^25^, gp100/IMDQVPFSV^26^, SSX-2/KASEKIFYV^27^, gp100/YLEPGPVTA^26^. For this, the T cells were knocked in with a pool of all 5 homology-directed repair templates (HDRTs) and stimulated with a pool of 5 different Ag-scaffolds carrying one specificity each (**Figure 4A**).

**Figure 4.**
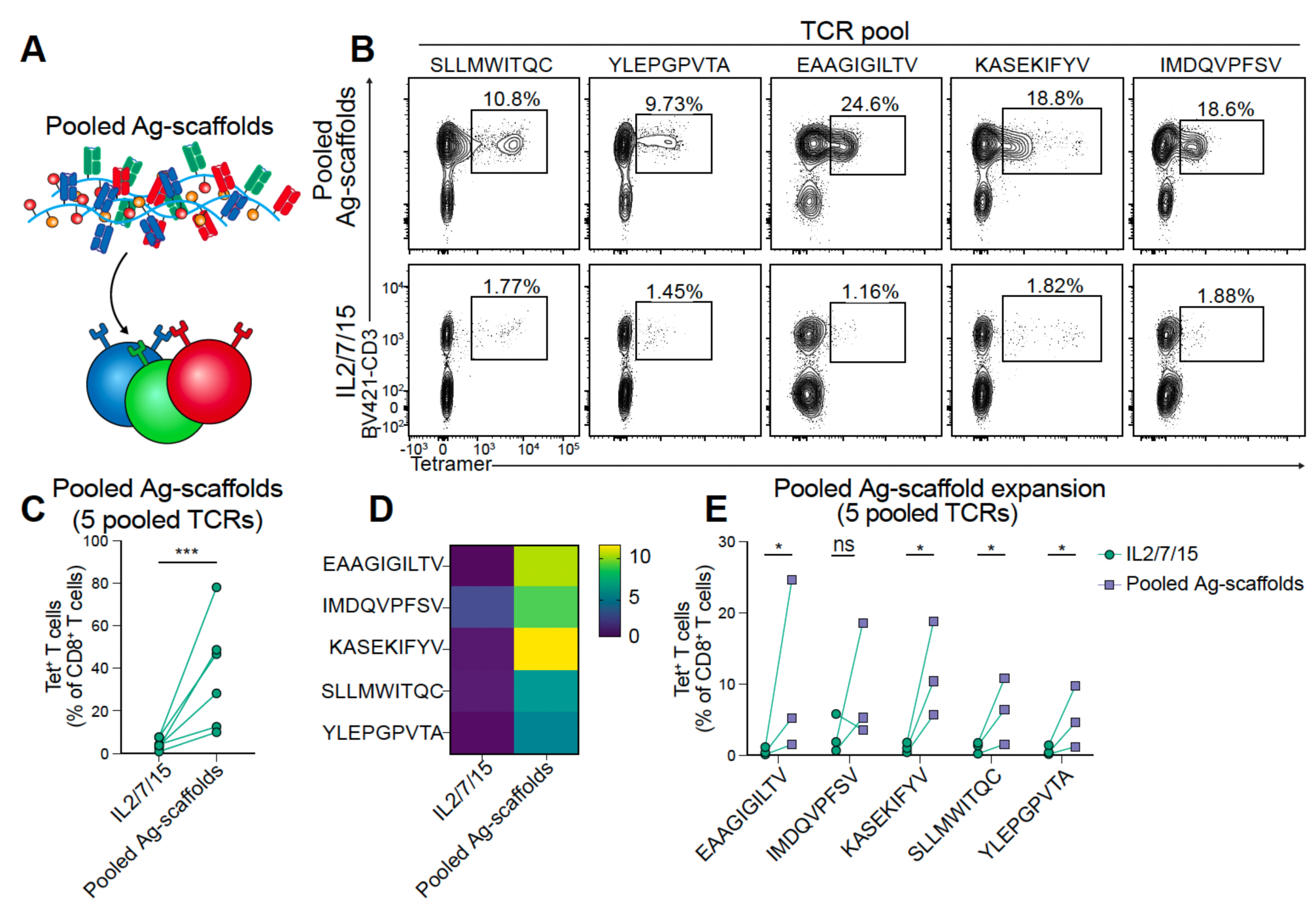
Pooled Ag-scaffold expansion of T cells engineered with a pool of TCRs. (**A**) Schematic depicting pooled Ag-scaffold expansion of a pool of T cells engineered to express 5 classical tumour shared antigen-specific HLA-A*02:01-restricted TCRs using CRISPR/Cas9 and representative flow plots showing percentages of Tet^+^ CD8^+^ T cells after 10 days of expansion using the pooled Ag-scaffolds or IL2/7/15. (**B**) Graph showing the total percentage of pooled tet^+^ CD8^+^ T cells after expansion (n=6 biological replicates from 2 experiments). (**C**) Heatmap showing the mean percentage of individual TCR Tet^+^ CD8^+^ T cells after expansion (n = 3). (**D**) Graph showing the % fold increase for each of the 5 KI TCRs in the culture following expansion with IL2/7/15 or Ag-scaffolds (n=3 biological replicates). Data in (**B**) is pooled from 2 experiments, and data in (**C, D**) is data from one experiment presented as dots for each data point in (**B, D**) and heatmap in (**C**). Statistical significance is evaluated by a ratio-paired T test in (**C**) and multiple ratio-paired t-tests with Holm-Šídák multiple comparison correction (**E**). P-values: *P < 0.05, **P < 0.01, ***P < 0.001, ****P < 0.0001.

Following 10 days of pooled expansion, we evaluated the expansion of the TCRs by staining using tetramers with separate fluorochromes per specificity. We found that the pooled expansions effectively expanded all 5 TCRs in the pool compared to the IL2/7/15 cytokine control from around 1-2 % to 10-25 % of CD8^+^ T cells **(Figure 4B).** When comparing the sum of all 5 specificities in cytokine and Ag-scaffold conditions, the pooled expansions resulted in a significant enrichment of the pool of antigen-specific T cells in the cultures **(Figure 4C).** It was clear that if the KI percentages were high (cytokine control), Ag-scaffold expansion would achieve higher final percentages in the cultures **(Figure 4C).** We observed specific expansion for all 5 TCRs in the pool **(Figures 4D-E)** with significant expansion observed for 4/5 TCRs in the pool **(Figure 4E).** Collectively, this data shows that the Ag-scaffold technology can be used to specifically expand a pool of TCRs following CRISPR/Cas9 KI.

### Ag-scaffold-expanded engineered T cells are highly cytotoxic *in vitro* and *in vivo*

To evaluate cytotoxicity and functionality of Ag-scaffold-expanded TCR and CAR T cells, CRISPR/Cas9 KI and lentiviral-modified TCR or CAR T cells were co-incubated with mCherry^+^ target cells and imaged on the Incucyte platform **(Figure 5A)**. CAR T cells were subjected to 3 rechallenges with mCherry^+^ CD19^+^ Jeko-1 cells to evaluate cytotoxicity, whereas TCR T cells were subjected to a single challenge with mCherry^+^ NY-ESO-1^+^/HLA-A*02:01^+^ A375 melanoma cells, and mCherry was imaged over 70 hours. First, we evaluated target cell killing following NY-ESO-1-specific TCR T cells in coculture using a single challenge with mCherry^+^ A375 cells. We found that Ag-scaffold-expanded NY-ESO-1-specific TCR T cells of both lentiviral and CRISPR/Cas9 origin were more effective at killing the target cell line than cytokine-expanded cells **(Figure 5A-C)**.

**Figure 5.**
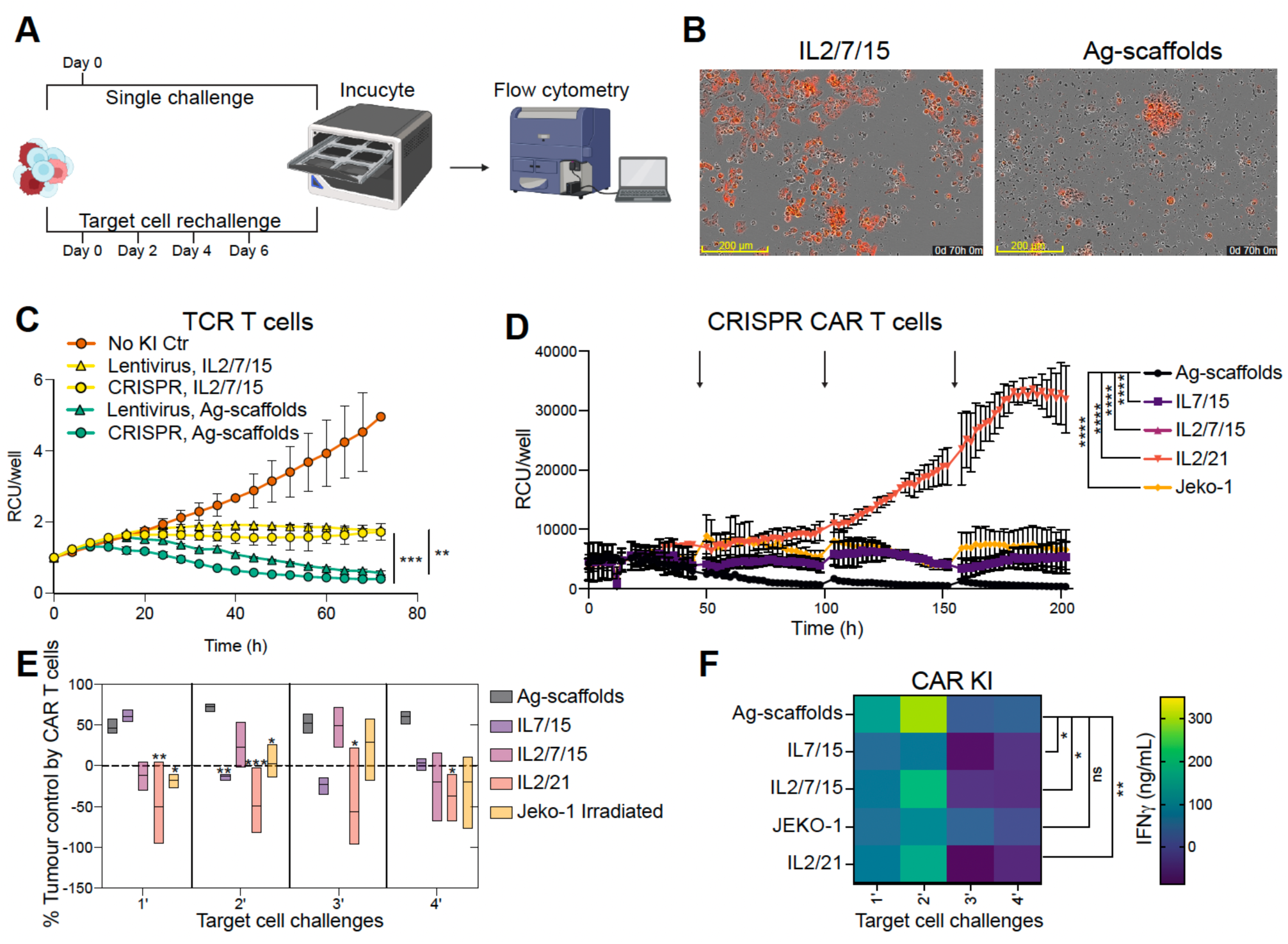
Antigen scaffold-mediated expansion yields CAR T cells that have high proliferative capacity, cytotoxicity, and cytokine production. (**A**) Schematic illustration of the Incucyte-based cytotoxicity re-challenge assay. (**B**) Representative images of mCherry^+^ A375 cancer cells after 70 hrs of coculture with cytokine-expanded (left) or Ag-scaffold-expanded (right) 1G4^+^ TCR T cells. Scale bar is 200 µm. (**C**) Cytotoxicity of CRISPR/Cas9 KI & Lentiviral 1G4^+^ TCR T cells in co-culture with mCherry^+^ A375 cells at an effector:target ratio of 1:1, tracked over 72 hours in the Incucyte, represented by relative confluence units (RCU) per well (representative of n=3 biological replicates). (**D**) Rechallenge cytotoxicity assay with CRISPR/Cas9 KI CAR T cells expanded with Ag-scaffolds, IL7/15, IL2/7/15, IL2/21, or Jeko-1 feeder cells mixed with mCherry^+^ Jeko-1 cells at a 1:1 effector:target ratio. CAR T cells were rechallenged with fresh Jeko-1 cells every 2 days (indicated by arrows), and cells were imaged for 200 hrs in the Incucyte, represented by relative confluence units (RCU) per well (representative of n=3 biological replicates). (**E**) Graph showing percentage tumour control by the CAR T cells over different target cell challenges calculated by %Tumour Control = (RCU at 48 hr – RCU at 0 hr)/(RCU at 0 hr)*100 (n=3 biological replicates from 1 experiment). (**F**) IFN-γ expression measured by ELISA in the supernatant (ng/mL) by CRISPR/Cas9 KI and lentiviral CAR T cells after every 48 hrs of target cell challenge as described in (**D**, **E**). (**C**-**F**) Statistical significance was determined using ordinary one-way ANOVA with Šídák’s multiple comparisons. In (**C**) the multiple comparison is comparing cytokine with Ag-scaffold for lentiviral and CRISPR/Cas9 conditions. In (**D**), the multiple comparisons compare Ag-scaffold to other groups. In (**E**) Ag-scaffold conditions were compared to all other groups within each timepoint by multiple comparisons (indicated with asterisks above the compared groups when P<0.05). In (**F**), the multiple comparisons compare the groups in the last rechallenge. P-values: *P < 0.05, **P < 0.01, ***P < 0.001, ****P < 0.0001.

Secondly, we evaluated CD19 CAR T cell cytotoxicity in a rechallenge assay with Jeko-1 cells as target cells, with rechallenges every 48 hours to simulate *in vivo* challenges. We found that Ag-scaffold-expanded CAR T along with irradiated Jeko-1- and IL7/15-expanded cells exhibited superior tumour control and cytotoxicity over the first two rounds of target cell rechallenges. However, Ag-scaffold-expanded CAR T exhibited significantly higher tumour control against CD19^hi^ Jeko-1 cells across rechallenges **(Figures 5A, D, E,** and **Figures S6A, B, C).** Normalised tumour control calculations revealed significantly higher tumour suppression by Ag-scaffold-expanded CAR T cells compared to those expanded with conventional cytokine cocktails.

Supernatants collected every 48 hours before each rechallenge revealed that Ag-scaffold-expanded CAR T cells secreted significantly higher levels of IFN-γ throughout all rechallenges than cytokine-expanded cells **(Figure 5F, Figure S6B).** Further, following rechallenges we evaluated the T cells by flow cytometry and observed that the Ag-scaffold-expanded CAR T cells had higher proportions of T_CM_ cells and lower percentages of terminally differentiated CAR T cells **(Figures 5C-F, Figures S6D-G)**

Finally, we tested the *in vivo* tumour-killing capacity of the CRISPR-engineered CD19 CAR T cells expanded with Ag-scaffolds or IL2/7/15 in NSG mice pre-inoculated with Jeko-1 lymphoma cells engineered to express luciferase (luc-mCherry^+^), allowing for tracking of tumour growth by bioluminescence (BLI). We evaluated a low dose of 0.25 and 0.5×10^6^ CAR T cells administered intravenously 7 days post inoculation to investigate the functional difference between the cytokine-expanded and Ag-scaffold-expanded CAR T cells. The mice were scanned using IVIS to monitor the change in BLI every 7 days after ACT. We found that treatment with CAR-T cells expanded with Ag-scaffolds resulted in complete tumour control in both the 0.25×10^6^ and 0.5×10^6^ CAR T cell groups until day 33 post treatment **(Figures 6A-B)**. The IL2/7/15 cytokine-expanded CAR T cells showed no significant tumour control with the lowest dose of 0.25×10^6^ CAR T cells, when compared to the no KI T cells, leading to the euthanisation of 4/5 mice by day 21 **(Figure 6B)**. The higher dose of 0.5×10^6^ cytokine-expanded CAR T cells slowed tumour growth until day 21 after which all mice had a significantly higher tumour burden when compared to the mice treated with Ag-scaffold-expanded CAR T cells **(Figure 6B)**. The mice were euthanised on day 33 and spleen and bone marrow were collected for the detection of CAR T cells by flow cytometry. We found that Ag-scaffold-expanded CAR T cells were present at significantly higher percentages and numbers in the spleen, but not in the bone marrow, compared to the cytokine-expanded T cells **(Figures 6C-D)**.

**Figure 6.**
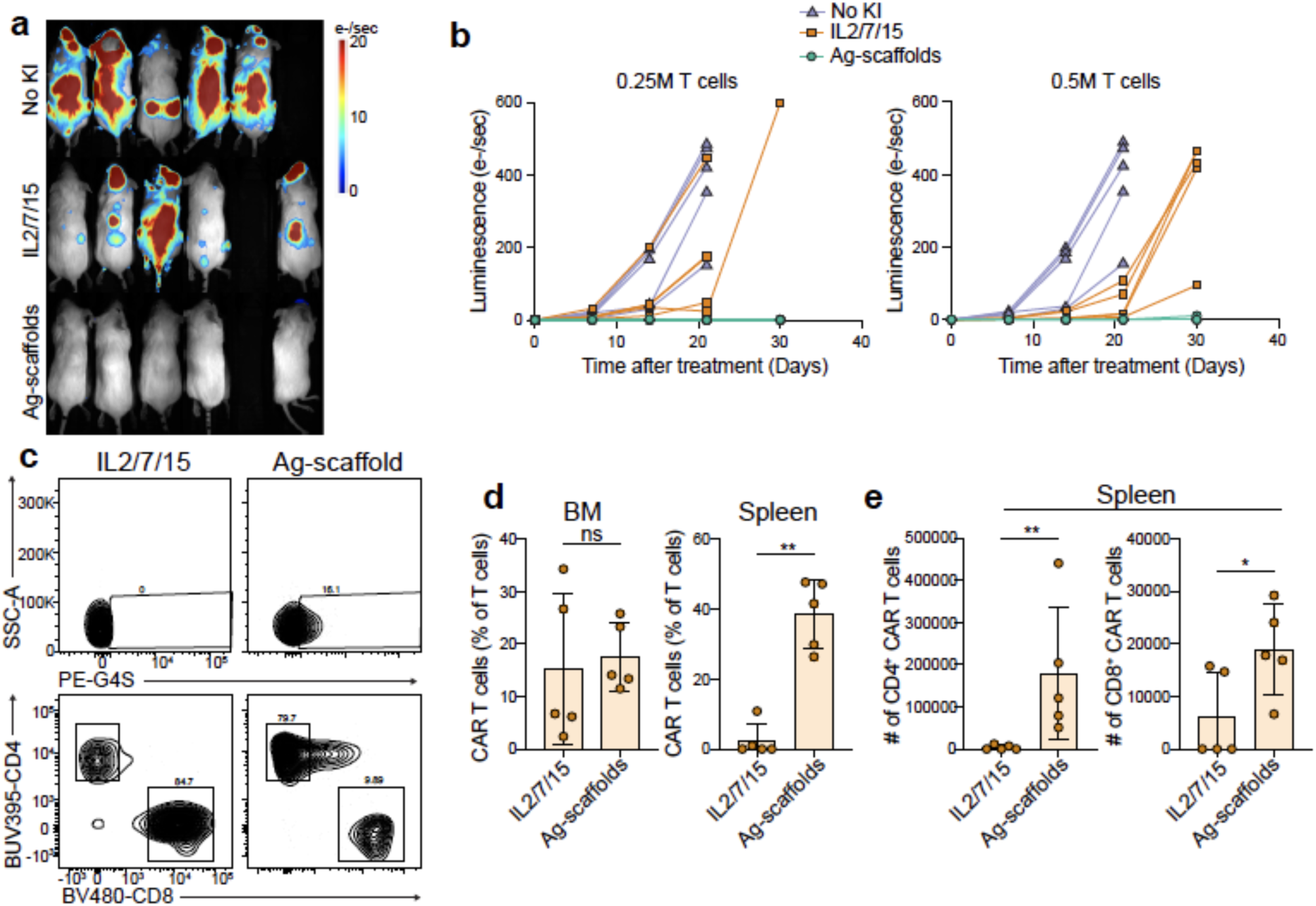
Antigen-scaffold-expanded CAR T cells are effective killers *in vivo* in NXG xenograft models. (**A**) BLI luminescence quantification of luciferase signal day 21 after intravenous inoculation of Luciferase-transduced JEKO-1 cell in immunodeficient NXG mice (n=5 mice per group) and injection of 0.5×10^6^ CAR T cells expanded with IL2/7/15 or Ag-scaffolds on day 7. Cell numbers were adjusted for the actual number of CAR-expressing T cells. **(B)** Quantification of cancer cell growth from luminescence signal (e-/sec) over time following injection of 0.25×10^6^ or 0.5×10^6^ CAR T cells 7 days after jeko-1 injection (n=5 mice per group). Mice were terminated when a predefined humane endpoint was reached. **(C-E)** On day 30, spleens and bone marrow (BM) from the IL2/7/15 and Ag-scaffold-expanded 0.5 ×10^6^ CAR T cell groups were analysed by flow cytometry, and CAR T cell percentages. (**C**) Representative flow plots showing CD3^+^ CAR T cell population (top) and CD3^+^CD4^+^ & CD3^+^CD8^+^ T cells (bottom), and number of CAR T cells (**D**) were quantified. Statistical significance in (**C, D**) determined by ratio-paired t-test: *P < 0.05, **P < 0.01, ***P < 0.001, ****P < 0.0001.

## Discussion

The substantial clinical efficacy of CAR T-cell therapy in treating haematological malignancies has brought significant attention to engineered T-cell therapies. However, the clinical production of these therapies faces several bottlenecks often resulting in final products with T cells of poor persistence and impaired function leading to suboptimal efficacy^11^. One major contributing factor to the compromised quality of T-cell products is the intense initial activation induced by anti-CD3 and anti-CD28 antibodies, followed by stimulation from recombinant cytokines. These cytokines are typically added in high concentrations to achieve clinically relevant numbers of specific T cells, often leading to T-cell exhaustion^28^.

The renewal and persistence of T-cell products are closely tied to the type and strength of co-stimulation and activation. An alternative to activation through CD3/CD28 cross-linking involves the use of irradiated target cells expressing the CAR antigen as feeder cells, which supports the expansion of antigen-specific T cells^29^. However, CAR products generated by this method show no significant functional improvement compared to those produced using conventional protocols and the co-stimulatory signals provided to the T cells can be difficult to tailor to a particular phenotype. Furthermore, the potential risk of re-infusing viable target cells into patients complicates the cGMP production of these products^30^.

Recently, a wave of innovation in bio-compatible and degradable materials for micro- and nanoscale aAPCs designed to expand antigen-specific T cells has emerged^18,31^. These aAPCs offer the advantage of creating customisable, off-the-shelf products targeting multiple specificities of TCRs and CARs. Additionally, their small size provides a spatial advantage by enabling paracrine delivery of cytokines and co-stimulation directly to antigen-specific cells upon engagement.

We recently developed a novel dextran scaffold-based technology and described how it is an effective tool for expanding endogenous viral and tumour antigen-specific CD8^+^ T cells^20^. Our previous research demonstrated the modularity of the streptavidin backbone, which facilitates the attachment of pMHC, and costimulatory cytokines such as IL2 and IL21 used in this study^21^. Building on this foundation, we hypothesised that this technology could be extended to enhance CAR and TCR T cell products. Especially the relatively low KI efficacy of CRISPR/Cas9-engineered T cells highlights the need to enrich engineered antigen-specific T cells without compromising their phenotype and functionality^6^.

Our results show that the average KI percentage is >15% in both TCR (∼15%) and CAR (∼6%) using Cas9 RNPs with double-stranded DNA HDR templates. The NY-ESO-1/HLA-A*02:01 pMHC Ag-scaffold supported expansion of NY-ESO-1 CRISPR/Cas9 TCR T cells, leading to a T-cell product with >90% TCR T cells. Similarly, the CD19 CRISPR/Cas9 CAR T cells expanded with CD19 CAR Ag-scaffold led to a ∼400-fold specific expansion of CAR T cells, reaching >60% CAR T cells on day 10 of expansion. Both yield a T-cell product that is significantly enriched compared to their cytokine-cocktail- and irradiated target cell-expanded counterparts. The strategy is likely also applicable to other receptor modalities, such as T cells engineered to present *de novo*-designed pMHC minibinder as antigen receptors^32–34^.

Ag-scaffold-expanded lentiviral and CRISPR/Cas9 TCR T cells exhibited robust proliferation while maintaining an efficacious early effector phenotype. These cells showed high expression of CD27 and CD28, markers associated with enhanced proliferative capacity and effector function, in contrast to cells expanded with IL2/7/15. The CD27^+^CD28^+^ phenotype has been correlated with improved disease prognosis in cancer immunotherapy^7,10,35^. Consequently, the Ag-scaffold-expanded 1G4 TCR T cells showed superior functionality in *in vitro* cytotoxicity assays in comparison to the IL2/7/15-expanded TCR T cells.

Ag-scaffold-expanded CAR T cells also showed remarkable functionality. Upon antigen re-stimulation, Ag-scaffold-expanded CAR T cells exhibited high proliferative capacity. These cells exhibited sustained tumour cell killing over four rounds of rechallenge, whereas CAR T cells expanded using conventional cytokines lost their cytotoxic capacity after two rounds of rechallenge. Moreover, Ag-scaffold-expanded CAR T cells consistently produced higher levels of IFN-γ during rechallenge assays.

Upon re-encountering antigen, previously primed T cells integrate signals from the TCR, co-stimulatory receptors, and cytokines to determine their fate: balanced stimulation drives further activation and differentiation into memory subsets (such as T_CM_ or T_SCM_), whereas excessive or prolonged signalling triggers activation-induced cell death (AICD)^36^. Serial antigen stimulation leads to T-cell exhaustion with high expression of markers of exhaustion^36,37^. We evaluated the efficacy of the Ag-scaffold-expanded CAR T cells in a relevant B-cell lymphoma xenograft mouse model. Even when the number of antigen-specific cells was normalised, the Ag-scaffold-expanded cells still outperformed the cytokine-expanded cells in clearing cancer cells. This strongly indicates that the Ag-scaffold-expansion drives a favourable phenotype, in addition to enriching the antigen-specific proportion of the cell product.

A significant limitation of TCR and CAR T-cell therapies for solid tumours is the heterogeneity of tumour antigen expression. Addressing this challenge often requires engineering T-cell pools with multiple specificities, but the bottleneck of low KI efficacy remains a barrier to producing clinically relevant products^4^. We demonstrated that scaffold technology could effectively enrich T cells engineered with different specificities, paving the way for customisable and efficient production of multi-specific T-cell products.

In conclusion, antigen- and cytokine-mediated activation and expansion using scaffold technology yielded highly enriched CAR and TCR T-cell products with superior potency *in vitro* and *in vivo* compared to conventional methods. These Ag-scaffold-expanded T cells demonstrated enhanced proliferation, sustained cytotoxicity, and improved cytokine production; *in vivo* superiority was established in a CD19^+^ lymphoma xenograft, and generalisation to TCR-engineered products and additional tumour models will be determined in future studies.

## Limitations of the study

While our study demonstrates that Ag-scaffolds can drive the preferential expansion of functional, genetically engineered T cells, several limitations warrant consideration. First, the *in vivo* efficacy of Ag-scaffold-expanded T cells was demonstrated exclusively using the CD19 CAR T cell model in a B-cell lymphoma xenograft. Although we established the *in vitro* cytotoxicity and phenotypic superiority of Ag-scaffold-expanded TCR T cells, the translational potential of the TCR-engineered products remains to be validated in relevant solid tumour animal models. Future studies will also have to evaluate safety of Ag-scaffold-expanded cells. Second, the starting material for all experiments consisted of peripheral blood mononuclear cells (PBMCs) obtained from healthy donors. T cells derived from heavily pre-treated cancer patients often exhibit compromised fitness and distinct baseline exhaustion profiles compared to healthy donors. Consequently, the expansion efficiency and functional rejuvenation observed here require validation using patient-derived material to fully ascertain clinical applicability.

## Resource availability

### Lead contact

Further information and requests for resources and reagents should be directed to the lead contact Sine Reker Hadrup.

### Materials availability

All reagents/materials generated in this study will be made available upon request. The request may require a completed Material Transfer Agreement.

### Data and code availability

Single-cell RNA-seq data have been deposited at the Gene Expression Omnibus (GEO) and are publicly available as of the date of publication. Accession numbers are listed in the Key Resources Table. All code used is publicly available as described in STAR Methods. Any additional information required to reanalyse the data reported in this paper is available from the lead contact upon request.

## Acknowledgements

We thank Sébastien Sebbaha from the Flow Cytometry and Imaging Core (FLIC) at DTU Health Tech, for flow cytometry experiments. We thank technicians Thi Minh Lien Nguyen and Panchale Olsen for assistance with cell cultures and ELISA.

The project was funded partly by a postdoctoral Lundbeck Foundation fellowship [R347-518 2020-2174] and the Novo Nordisk Foundation (Challenge program, Nano-Immune-Cell-Engineering, NICE) [NNF21OC0066562], European Union’s Horizon 2020 research and innovation program Marie Curie international training network, T-OP [955575].

## Author Contributions

Shared authors have contributed equally and are listed alphabetically by last name, and they are free to change the order on their CVs. S.R.H., K.H.J., K.R., and M.O. conceived the project and provided funding. S.R.H. and M.O. supervised the work. K.R., K.H.J., S.R.H., and M.O. wrote and reviewed the manuscript. K.H.J., K.R., C.R.P., A.K.G., L.F.V., H.R.H., S.T., L.S.G., E.D., A.H.S., L.P.F., M.B., and M.S.F. performed experimentation and analysis for the manuscript. All authors approved of the manuscript.

## Materials and Methods

### Key Resources table

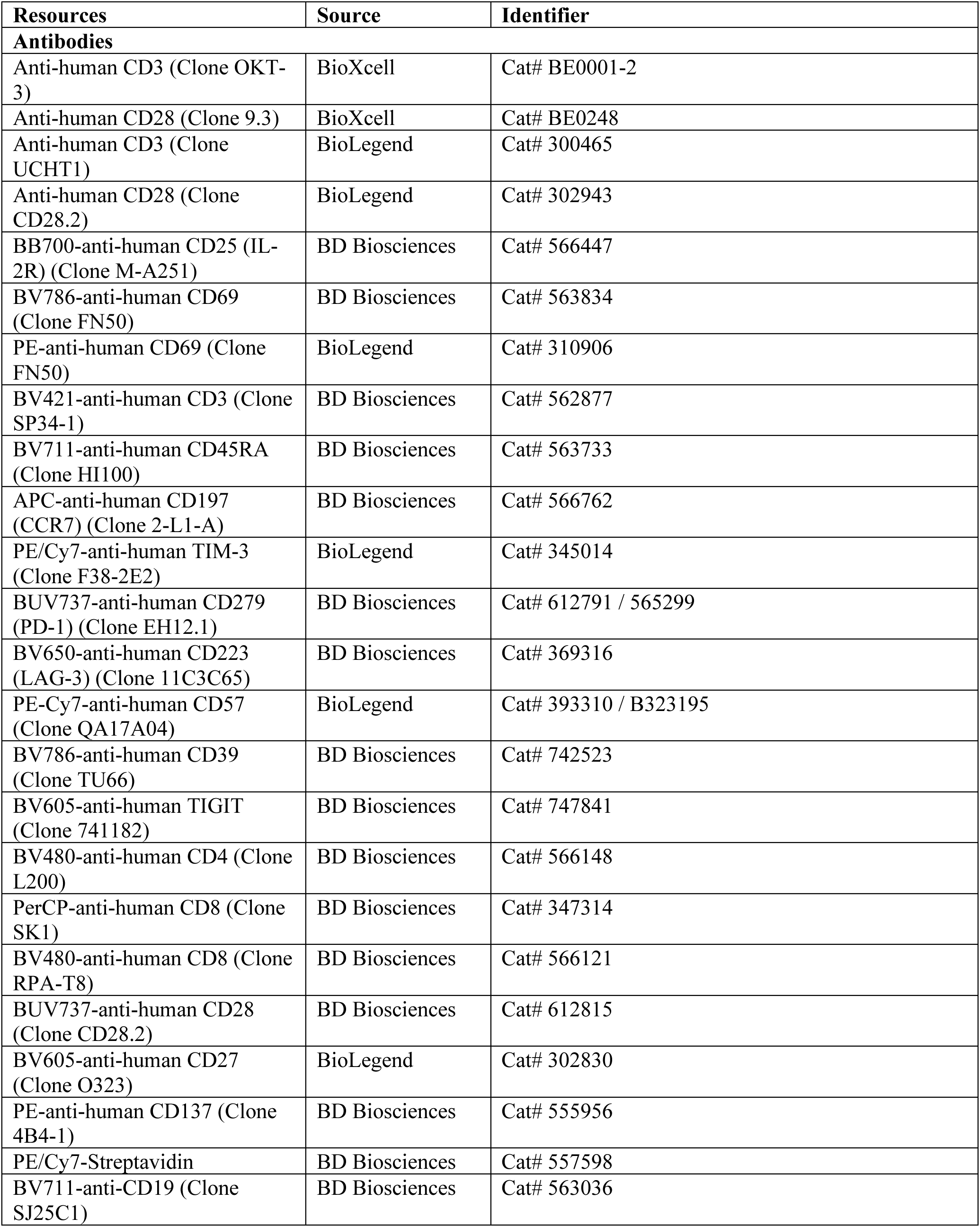
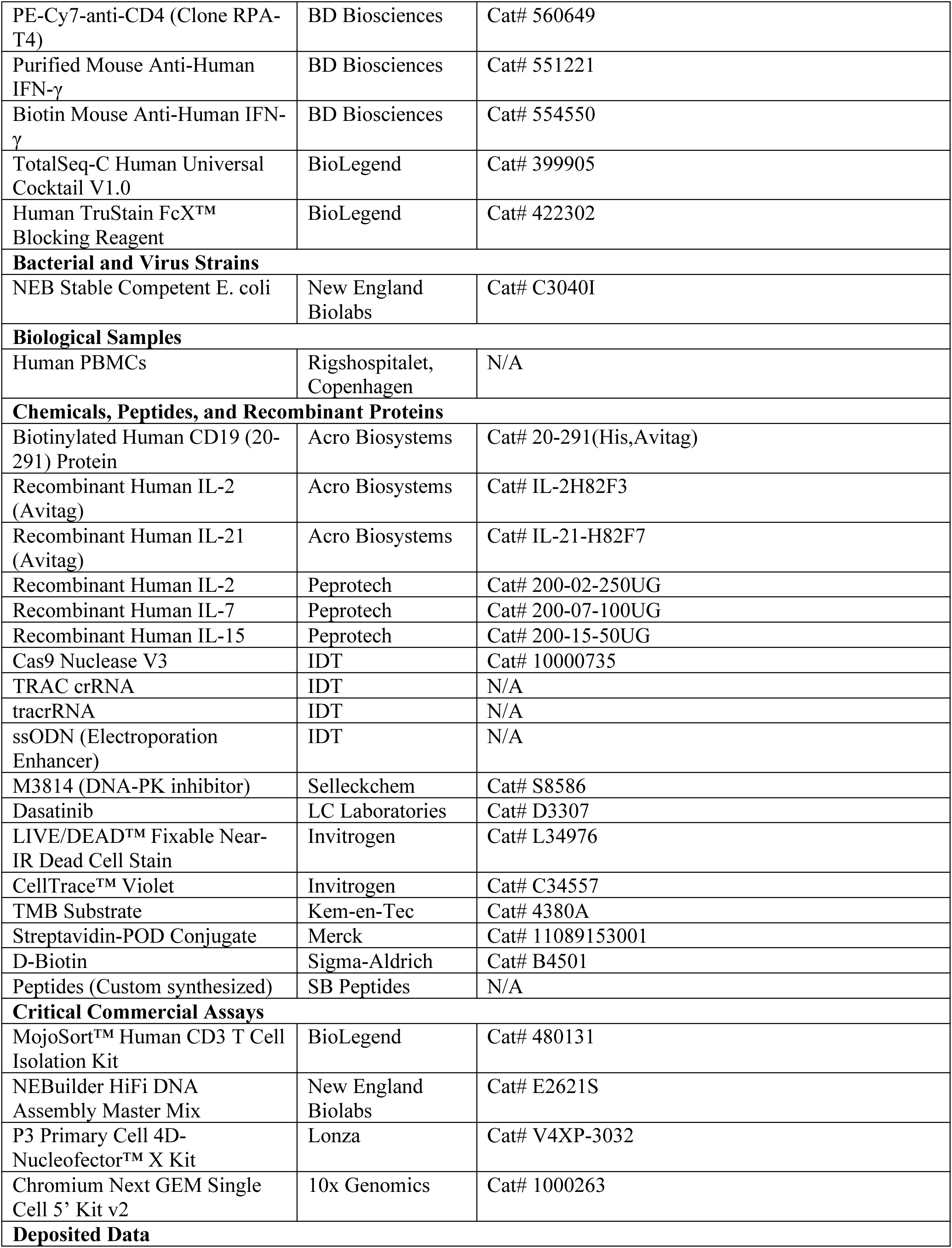
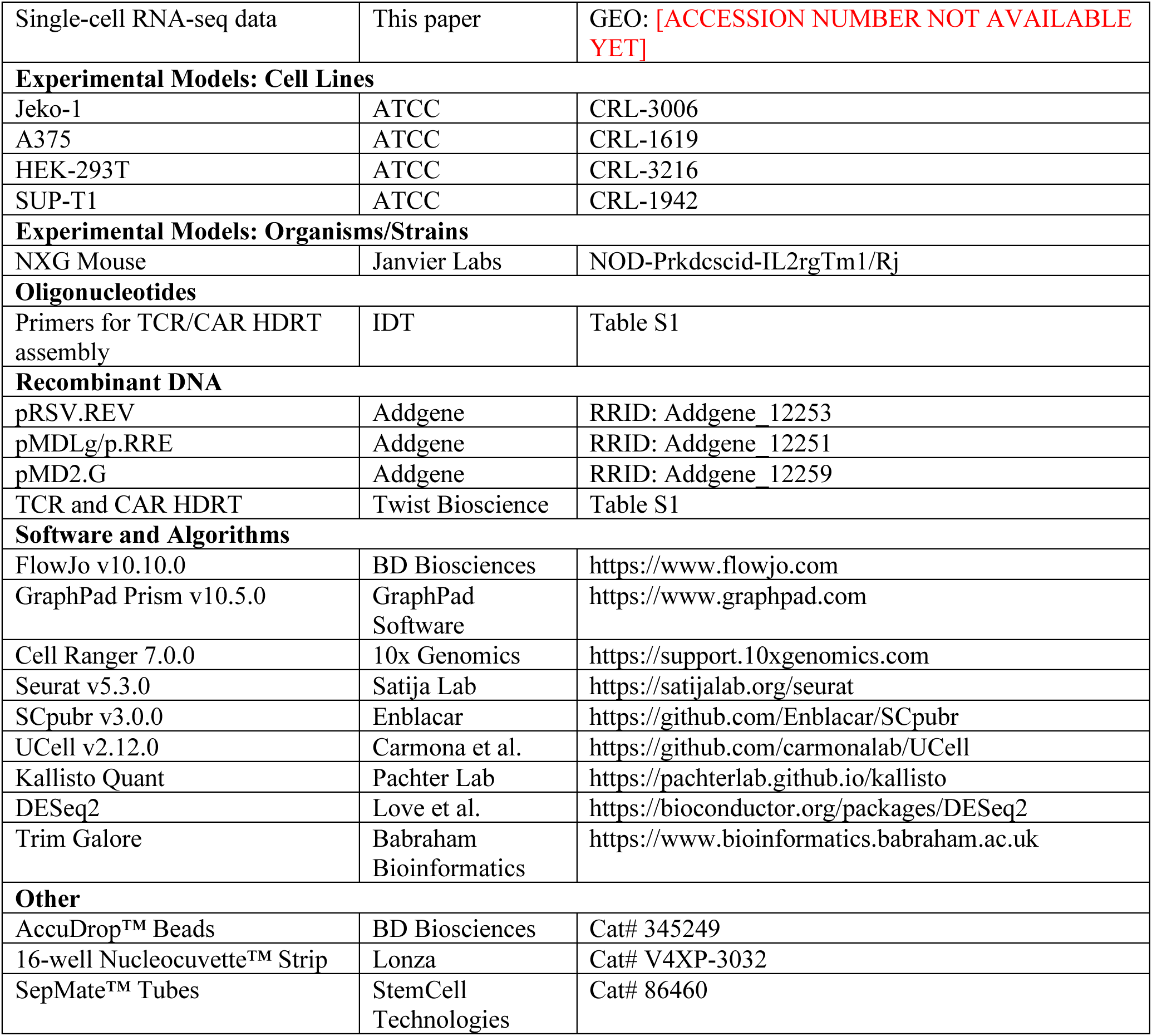

### Experimental model and study participant details

#### PBMCs

PBMCs were sourced from healthy donors from Rigshospitalet, Copenhagen, approved by the Danish ethics committee. PBMCs were isolated from whole blood by density centrifugation using SepMate tubes (StemCell) and frozen in liquid nitrogen in FBS and 10 % DMSO.

#### T cell cultures

Cryopreserved PBMCs were thawed and washed in RPMI-1640 GlutaMAX™ + 10 % FBS (R10 medium), and T cells were isolated by negative selection using MojoSort™ Human CD3 T Cell Isolation Kit (Biolegend) according to the manufacturer’s instructions. Cells were counted and activated on 5 µg/mL plate-bound anti-CD3 (clone: UCHT1, Biolegend) with 2 µg/mL soluble anti-CD28 (clone: 28.2, Biolegend) in X-vivo-15 (Lonza, BE02-060Q) + 5 % human serum (Sigma Aldrich Heat Inactivated H3667), supplemented with 200 IU/mL recombinant human IL2 (Peprotech), 5 ng/ml IL7 (Peprotech), and 5 ng/ml IL15 (Peprotech).

#### Cell lines and culture

The human cell lines Jeko-1 (B cell lymphoma) and A375 (melanoma) were obtained from American Type Culture Collection (ATCC) and cultured in RPMI supplemented with 10% fetal bovine serum and 1% penicillin-streptomycin.

For cytotoxicity assay and *in vivo* tumour models, these cell lines were transduced with mCherry-Luc lentiviral vector (**Table S1**) and were sorted on a FACSmelody (BD Biosciences) to establish new cultures.

#### In vivo NXG tumour xenograft models

The Danish National Animal Experiment Inspectorate and the institutional ethics board at the Technical University of Denmark approved all procedures (License#2020-15-0201-00748). For the efficacy study, 7–10-week-old female immunodeficient NXG mice (NOD-Prkdcscid-IL2rgTm1/Rj; Janvier Labs) were intravenously injected with 1 × 10⁶ luciferase-expressing Jeko-1 cells. After a 7-day tumour engraftment period, mice were randomised and injected intravenously with either the Ag-scaffold- or IL2/7/15-expanded CD19 CAR T cells in 200 µL PBS. Effector:target ratios were adjusted to the CAR-positive fraction, i.e., the number of cells shown is the number of CAR^+^ T cells. Tumour burden was monitored weekly by bioluminescence imaging (Optical Imaging unit, MILabs). For imaging, mice were injected intraperitoneally (i.p.) with D-Luciferin (3 mg/mouse) and imaged 20 minutes later. Luminescence was then quantified using fixed-size regions of interest. Body weight and general health were monitored throughout the study. At termination, single-cell suspensions of spleen and bone marrow were prepared for flow cytometry. Spleens were processed through a 70 µm filter, followed by red blood cell lysis. Cells were then stained with a viability dye (Near-IR) and antibodies for CD3, CD4, CD8, CD45RA, CCR7, and a CD19 CAR tetramer before analysis on an LSRFortessa™.

### Method details

#### Preparation of dsDNA homology-directed repair template

The HDRTs used for KI of the NY-ESO-1-specific 1G4 TCR were designed as follows: 5’ HA (300 bp)-T2A-TCRb-furin cleavage site-SGSG linker-P2A-TCRa variable chain-3’ HA (300 bp)^4,38^, whereas HDRTs used for KI of CARs were designed as follows: 5’ HA (300 bp)-T2A-scFv-CD8 hinge/transmembrane domain-41BB/CD3z signalling domain-bGH polyA-3’ HA (300 bp) (Table S1). For the four other TCRs, a 500 bp 3’ and 5’ HA was used. TCR constructs were ordered as clonal genes (Twist Bioscience), CAR constructs were assembled by HiFi assembly (NEB, E2621S). Briefly, we performed PCR of the backbone from the TCR HDR plasmid containing the TRAC HDR homology arms (primers: TCR_HDR_HiFi_fwd, TCR_HDR_HiFi_rev), the CD19 CAR gene from the lenti_CD19_CAR plasmid (primers: CD19 plasmid_fwd, CD19 plasmid_rev), and a bGH polyA tail (primers: pA AAVS1 plasmid_fwd, pA AAVS1 plasmid_rev) and purified using a PCR clean-up kit (Qiagen, #28104). The PCR fragments were mixed in a HiFi reaction according to the manufacturer’s instructions, followed by transformation in NEB Stable bacteria (NEB, #C3040I). Assembled plasmids were sequence verified. The HDRTs were amplified from the plasmids using TRAC_HDRT_1.Fwd and TRAC_HDRT_1.Rev primers (**Table S1**) with Herculase Fusion II polymerase with the following PCR program: 1 cycle of 95°C, 5 min. 34 cycles of 95°C, 20 sec; 63°C, 20 sec; 72°C, 1.5 min. 1 cycle of 72°C, 3 min. TRAC_HDRT_2.Fwd and TRAC_HDRT_2.Rev were used for 500 bp HAs. Products were purified using 1:1 volume homemade Ampure XP beads^39^. Concentration of dsDNA HDRT was measured by Nanodrop, and products were validated by gel electrophoresis.

#### Non-viral CRISPR/Cas9-mediated KI

Our non-viral CRISPR/Cas9-mediated KI protocol was adapted from previous publications^4,40^. Briefly, gRNAs were prepared by mixing equimolar ratios of 200 µM TRAC_crRNA and tracrRNA (IDT) resuspended in duplex buffer, and incubated at 95°C for 5 min or 37°C for 30 min and slowly cooled to RT. To prepare Cas9 RNPs, 3 µL of PBS, 1.2 µL of 100 µM gRNA, and 1 µL of Cas9 (62µM Cas9 nuclease V, IDT) were mixed and incubated at RT for 15 min. 0.5 µL of scrambled Single-stranded oligodeoxynucleotides was added as electroporation enhancer (ssODN, 200 µM, IDT). 1 µg of double-stranded HDRT was added 1 min before mixing with T cells for nucleofection.

1×10^6^ T cells per nucleofection cuvette were counted and spun down at 100xG for 10 min and resuspended in 20 µL RT P3 buffer (Lonza). RNP/HDRT mix was added, and cells were transferred to a 16-well nucleofection cuvette (Lonza) and nucleofected with pulse-code EH115 on a 4D-nucleofector. 80 µL of incubator equilibrated X-vivo-15 without serum or cytokines was added and cells were incubated for 30 min. Cells were transferred to a 48 well plate at 2×10^6^ cells/well in X-vivo-15 + 5 % HS + 200 IU/mL rhIL2, 5 ng/mL IL7, 5 ng/mL IL15, and DNA-PK inhibitor 1 µM of M3814 (S8586, Selleckchem). M3814 inhibitor was removed the next day. KI was evaluated 2-3 days after nucleofection.

#### Lentiviral transduction of T cells

The CAR/TCR was inserted into a third-generation lentiviral vector under the control of a human EF1α promoter. All plasmids were synthesised by GenScript, USA. Lentiviral particles were generated in HEK-293T cells by transfecting them with three separate packaging plasmids: pRSV.REV(Addgene plasmid #12253), pMDLg/p.RRE (Addgene plasmid #12251), and pMD2.G (Addgene plasmid #12259) were a gift from Didier Trono^41^, along with a transfer plasmid containing the DNA construct of interest. The titer of the produced lentivirus particles was assessed through a titration experiment by infecting SUP-T1 cells and subsequently analysing GFP or surface expression of the receptor via tetramer staining via flow cytometry.

The T cells were activated 2×10^6^ cells/mL for 48 hrs at 37°C and 5% CO_2_ in x-vivo-15 media (Lonza) supplemented with 5% human serum containing recombinant IL2 (200IU/mL), IL7 (5ng/mL), IL15 (5ng/ml) (Preprotech, USA) and anti-CD3 and CD-28 (5ng/ml). After the activation period, the T cells were washed and resuspended in x-vivo-15 media (Lonza) supplemented with 5% human serum and transduced with the required amount of lentivirus at a multiplicity of infection (MOI) of 1 and expanded for another 10 days.

#### Preparation of peptide MHC

pMHC monomers and pMHC tetramers were prepared as previously described^42,43^. Briefly, the peptides were synthesised and acquired from SB peptides and resuspended in DMSO at 10 mM, and in-house produced avitagged-HLA-A*02:01^Y84C^ was used for peptide loading (final conc.: 100µg/ml HLA; 200 µM peptide) at RT for 30 min prior to tetramerising with PE- or APC-labelled streptavidin for 30 min on ice and blocking with D-biotin 20 min on ice prior to freezing in 0.5% BSA and 5% glycerol at -20°C.

#### Antigen scaffold generation

Ag-scaffolds were assembled, using a streptavidin-conjugated dextran backbone (500 kDa) (Fina Biosolutions) mixed with biotinylated peptide MHC class I complex or CAR antigen (CD19 Avitag, Acro Biosystems), IL-2 Avitag (Acro Biosystems #IL-2H82F3), and IL-21 Avitag (Acro Biosystems #IL-21-H82F7), followed by incubation at 4°C for 30 min. Ag-scaffolds were incubated with D-Biotin at 4°C for 20 min to block potential free streptavidin sites on the dextran backbone. To remove any unbound molecules, Ag-scaffolds were filtered through 100 kDa spin columns (Vivaspin6, Sartorius). Ag-scaffolds were frozen in 0.5% BSA and 5% glycerol at −80°C for up to 15 months^21^.

#### Flow cytometry

Cells were stained with antibodies against phenotypic and intracellular markers, then run on a Fortessa (BD) and the data analysed using FlowJo software. For gating strategies (see **Figures S7A, B, and Figure S8A).** The antibodies used for surface and intracellular marker staining are listed in Key Resouce Table.

Cell lines were sorted using a FACSAria flow cytometer (BD, USA) with a 100 µm nozzle. Amplitude was adjusted to optimise droplet break-off, and droplet calibration was performed before each sort using AccuDrop™ beads (BD, USA).

#### Tetramer staining

Cells were centrifuged at 500 x g for 5 minutes at 4°C, and the supernatant was discarded. The pelleted cells were then resuspended in 5 µL of 1 µM Dasatinib (LC Laboratories, #D3307), antigen tetramers were added, and the cells were incubated for 15 minutes at 37°C in the dark. The cells were washed once with FACS buffer, followed by staining with Near-IR (NiR) viability dye (Invitrogen, #L34976) and additional antibodies for surface staining for surface markers 30 min at 4°C in the dark. Finally, the cells were washed twice with FACS buffer in 200-300 µL of FACS buffer and immediately analysed on a LSRFortessa flow cytometer (BD, USA). Alternatively, the cells were fixed with 50 µL of 1% paraformaldehyde for 1-2 hrs (if required), washed twice with FACS buffer, and analysed 2-24 hrs later^44^

#### Proliferation assay

The T cells were resuspended at 1×10^6^ cells/mL in PBS and 0.75µL CellTrace violet/mL of PBS was added and incubated at 37°C for 15 mins. After incubation, the cells were washed 2x and co-cultured with irradiated Jeko-1 target cells at a 1:5 effector-to-target ratio for 6 days. Flow cytometry was used to measure CellTrace Violet dilution, and the results were analysed using FlowJo software (BD, v10.10.0).

#### Incucyte

The CD19 CAR T cells and the CD19 target cells, Jeko 1 lymphoma cell line modified to express mCherry, resuspended in RPMI supplemented with 10%FBS and 1% PenStrep are seeded at the mentioned effector:target ratio. In Incucyte assays, effector:target ratios were adjusted to the CAR-positive fraction, e.g., 1:1 (10,000 target cells/well: 10,000 CAR^+^ T cells adjusted based on the % of CAR T cells present in the final expansion product) on day 0 in a 96-well plate. Plates were incubated at 37°C and 5%CO_2_. After 48 hrs of incubation, the supernatant is collected for ELISA, and the wells are rechallenged with 10,000 target cells in 200µL/well. This process is repeated every 48 hrs, to simulate the challenge that the CAR T cells will encounter *in vivo*.

The A375 target cells (mCherry-luciferase modified) expressing the target antigen were plated 5000 cells/well and incubated overnight, and the CRISPR/Cas9 or Lentivirus-engineered T cells expanded with either IL2/7/15 or Ag-scaffold were added on top at an effector target (E:T) ratio of 4:1 or 1:1 adjusted based on the % of TCR^+^ T cells present in the final expansion product.

#### Intracellular staining

The expanded TCR T cells were incubated with the A375 target cells in an effector:target ratio (E:T) of 1:1 for 6 hrs in RPMI media supplemented with αCD107s-PE antibody and GolgiPlug. Negative control was supplemented with media, and the positive control was incubated with a leukocyte-activating cocktail. The T cells were harvested and stained for dead cells and surface markers (αCD3, αCD8) as described previously and the cells were fixed with 4% paraformaldehyde and incubated for 30 mins on ice and permeabilised (Invitrogen, #88-8824-00) and stained for intracellular markers (TNFα and IFN-γ) for 30 minutes at 4°C in the dark (Key Resource Table). Finally, the cells were washed twice, resuspended in 200 µL FACS buffer, and analysed on an LSRFortessa.

#### ELISA

96-well plates are coated with purified mouse anti-human IFN-γ (BD, #551221) resuspended in 0.1 M Na_2_HPO_4_ buffer at (1µg/mL) and coated with 50 µL/well and incubated at 4°C overnight. The coating was removed, and the plates were blocked with blocking buffer (1% BSA in 1xPBS) for 2 hrs at RT on a plate shaker at 300 rpm. Blocking buffer was discarded, and samples and standards were incubated at 50 µL/well for 2 hrs at RT at 300 rpm. Following incubation, the plate was washed with PBS-Tween buffer and incubated with 50 µL/well (0.5 µg/mL) of the detecting antibody, Biotin mouse anti-human IFN-γ (BD, #554550), for 1 hr at RT at 300 rpm. The plate was washed with PBS-Tween, and streptavidin-POD conjugate (Merck, #11089153001) was added (50 µL/well) and incubated for 30 mins at RT at 300 rpm. Finally, the plate was washed with PBS-Tween, and 50 µL/well TMB (Kem-en-Tec, #4380A) solution was added (50 µL/well). The reaction was stopped by adding 50 µL of H_2_SO_4._ OD450 was measured. The analysis was done using the Dose-response curve.

#### Single-cell RNA sequencing

Expanded T cells were stained with multimers and antibodies as previously reported^45,46^. In brief, cells were washed twice with PBS+0.5% BSA. 3µL of the DNA-barcoded pMHC multimer (SLLMWITQC/HLA-A*02:01) was added to the T cells and stained at a final volume of 100 µL for 60 min on ice, followed by three washes with PBS+0.5% BSA. Cells were blocked with 2.5µL of Human TruStain FcX Blocking reagent (BioLegend, 422302) for 10 min on ice. TotalSeq-C Human Universal Cocktail V1.0 (BioLegend, 399905) was added and incubated for 10 min, whereafter TotalSeq-C anti-Human Hashtag antibodies (#1-#24) and fluorochrome-conjugated lineage antibodies (LIVE/DEAD™ Fixable Near-IR Dead Cell Stain Kit, BV421-anti-human CD3-BD Biosciences (562877), BV480-anti-human CD8-BD Biosciences(566121), PE-Cy7-anti-CD4-BD Biosciences (560649)) were added, followed by 30 min on ice. Finally, the cells were washed three times prior to acquisition. Cells were sorted into PBS+0.5% BSA and centrifuged at 390g at 4℃ for 10 min. The cells were resuspended in ∼20 µL of the supernatant and loaded into a Chromium Controller and prepared for sequencing as described in the Chromium Next GEM Single Cell 5’ v2 protocol (CG000330).

#### Single-cell RNA sequencing analysis

Gene expression (RNA), antibody-derived tag (ADT), and hashtag oligonucleotide (HTO) libraries were sequenced on an Illumina platform at Novogene. Reads were processed using CellRanger multi (cellranger-7.0.0, 10X genomics) with feature references associating barcodes to CITE-seq antibodies and hashing antibodies (reference transcriptome: GRCh38-1.2.0) and analysed using Seurat (V.5.3.0)^47^. Counts for hashtags were normalised and then demultiplexed using HTODemux and only cells identified as singlets were used for downstream analysis.

Quality control filtering was applied to remove low-quality cells^48^. Cells expressing <100 and >5,500 genes were removed, along with cells with an RNA count of >20,000. Furthermore, cells with >10% mitochondrial reads were removed. Following filtration, a total of 2,292 cells were included in further processing and analysis. The RNA data were normalised, and the effect of the cell cycle was removed by regressing out the S and G2/M scores during data scaling. A set of 2,000 highly variable genes was defined using the FindVariableFeatures function and subsequently utilised for dimensionality reduction via principal component analysis. Cell clustering was performed with the FindNeighbors and FindClusters functions, and Uniform Manifold Approximation and Projection (UMAP) plots were generated using the do_Dimplot function from the SCpubr (V.3.0.0) package^49^. Gene signatures were scored using the AddModuleScore_UCell function from UCell (V.2.12.0) with cytotoxicity being defined by the gene set GZMA, GZMB, GNLY and PRF1, dysfunctionality being defined by the gene set HAVCR2, LAG3, ENTPD1, TCF7-, SLAMF6*-*, and proliferation being defined by the gene set CCDN3, CDK6, MKI67, PCNA, MCM3, MCM5, MCM7, CLSPN, IL2RA^50^. Violin plots were generated using the VlnPlot function with modifications for adding statistics using the non-parametric Mann-Whitney U test.

#### Bulk RNA sequencing and differential gene expression analysis

Sorted CD3^+^CAR^+^ T cells, previously expanded using either Ag-scaffolds or cytokines, were submitted for bulk RNA sequencing at Genewiz*®*. The resulting raw reads were processed using Trim Galore, and transcript abundances were subsequently quantified with Kallisto Quant^51^. We performed differential gene expression analysis across three donors using DESeq2 in R^52^. Genes were considered differentially expressed if adjusted p-value < 0.05 and Log2 fold change > 2 (upregulated) or < -2 (downregulated). A volcano plot of the results was generated using ggplot2, and a heatmap of selected differentially expressed genes was created with ComplexHeatmap^53^.

### Quantification and statistical analysis

For flow cytometry analysis, FlowJo (v10.10.0) was used, and numerical data analysis was performed in GraphPad Prism (v10.5.0). Statistical analysis was performed for normally distributed paired data using ratio-paired T tests. When comparing more than two groups, we used an Ordinary One-Way ANOVA with the Holm-Šídák multiple-comparison correction, unless stated otherwise. Data is presented as individual data points, with mean±SD indicated and *P < 0.05, **P < 0.01, ***P < 0.001, ****P < 0.0001.

