## Supplementary material for "Antigen-scaffolds drive preferential expansion of functional genetically engineered CAR and TCR T cells": Figure S

**Supplemental material**

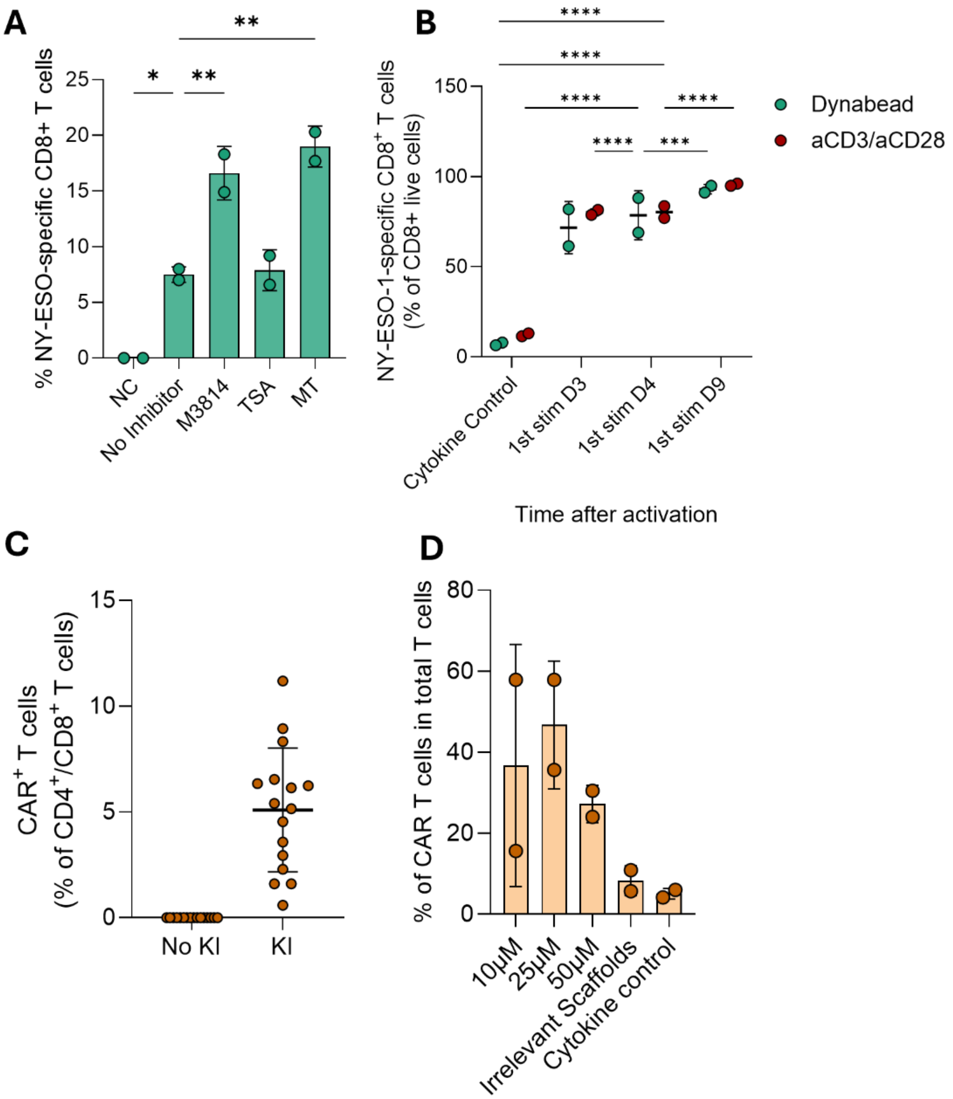


**Figure S1. Optimization of knockin of CAR and TCRs in primary T cells using CRISPR/Cas9.** (**A**) Graph showing % of TCR$^+$, CD8$^+$ T cells after 48 hrs of CRISPR/Cas9 TCR and incubation in different non-homologous end-joining inhibitors for 24 hrs. (**B**) Graph showing the fold expansion of TCR$^+$CD8$^+$ T cells 14 days post activation by Dynabeads or α-CD3/CD28 antibody for KI (12 days post TCR KI) with Ag-scaffolds added at different days after initial activation of T cells in comparison to T cells expanded with cytokine cocktail throughout the protocol. (**C**) Graph showing the KI efficacy of CD19 CAR HDRT in the T cells 48 hrs post-KI of the CD19 CAR HDRT in TRAC locus using CRISPR/Cas9. (**D**) Graph showing the percentage of CD19 CAR T cells expanded with media supplemented with 10, 25, or 50μM of CD19 antigen conjugated Ag-scaffolds, irrelevant scaffolds conjugated with CD79b antigen, or IL2/7/15 cytokine cocktail 14 days post-KI of the CD19 CAR HDRT in TRAC locus using CRISPR/Cas9. Data in

(**A**) and (**B**) represent results from one experiment, presented as mean ± SD. Statistical significance was determined by ordinary one-way ANOVA with Holm-Šídák's multiple comparisons test: *$P < 0.05$, **$P < 0.01$, ***$P < 0.001$, ****$P < 0.0001$.

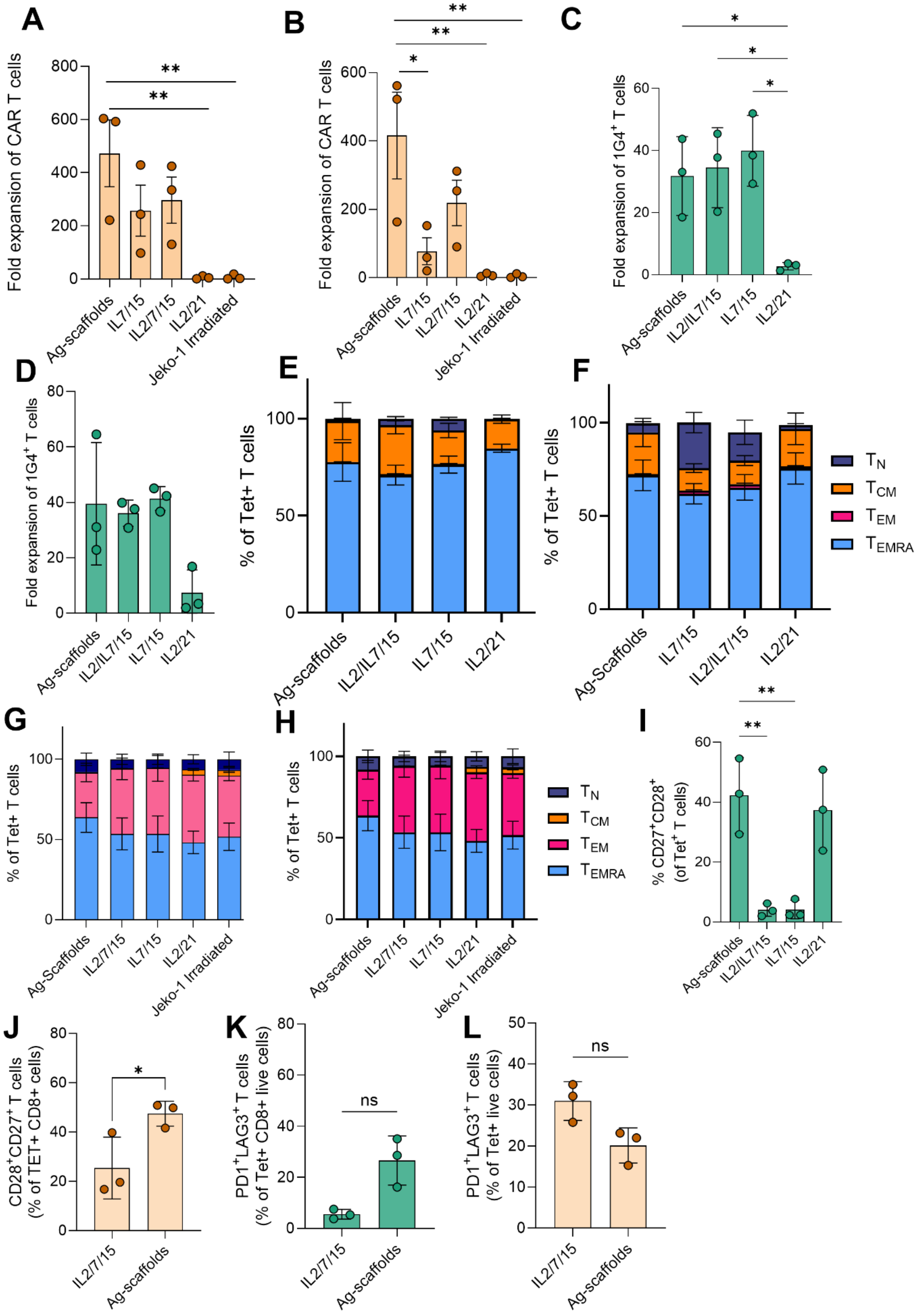

A
Fold expansion of CAR T cells
Ag-scaffolds
IL7/15
IL2/7/15
IL2/21
Jeko-1 Irradiated
B
Fold expansion of CAR T cells
C
Fold expansion of 1G4+ T cells
IL2/IL7/15
D
Fold expansion of 1G4+ T cells
E
% of Tet+ T cells
F
% of Tet+ T cells
Ag-Scaffolds
TN
TCM
TEM
TEMRA
G
% of Tet+ T cells
H
% of Tet+ T cells
I
% CD27+CD28+ (of Tet+ T cells)
J
CD28+CD27+ T cells (% of TET+ CD8+ cells)
K
PD1+LAG3+ T cells (% of Tet+ CD8+ live cells)
ns
L
PD1+LAG3+ T cells (% of Tet+ live cells)
ns

**Figure S2**. **Expansion and phenotypic analysis of CAR and TCR T cells. (A-B)** Fold expansion of CAR T cells over 10 days expansion following lentiviral transduction (**A**) or CRISPR/Cas9 KI (**B**). **(C-D)** Fold expansion of TCR T cells over 10 days following lentiviral transduction (**C**) or CRISPR/Cas9 KI (**D**). (**E-F**) Phenotypic characterization of CRISPR/Cas9 (**E**) and Lentiviral (**F**) CAR T cells, based on CCR7 and CD45RA expression, showing $T_N$ ($CD45RA^+CCR7^+$), $T_{CM}$ ($CD45RA^-CCR7^+$), $T_{EM}$, and $T_{EMRA}$ subsets. (**G-H**) Phenotypic characterization of CRISPR/Cas9 (**G**) and lentiviral (**H**) TCR T cells based on CCR7 and CD45RA expression, showing $T_N$ ($CD45RA^+CCR7^+$), $T_{CM}$ ($CD45RA^-CCR7^+$), $T_{EM}$, and $T_{EMRA}$ subsets. (**I**) Graph shows lentiviral TCR T cell percentage of $CD27^+CD28^+$ population of the total CAR T cells expanded with Ag-scaffolds or IL2/7/15. (**J**) Graph shows lentiviral CAR T cells percentage of $CD27^+CD28^+$ population of the total CAR T cells expanded with Ag-scaffolds or IL2/7/15. **(K-L)** Graph showing the percentage of lentiviral engineered $CD8^+$ TCR **(K)** and $Tet^+$ CAR T cells **(L)** expressing PD1 and LAG3 surface markers. Data in (**A, B, C, I**) represent pooled results from one experiment, presented as mean ± SD. Statistical significance was determined by One-Way ANOVA with Holm-Šídák's multiple comparisons test; (**J**) statistical significance was determined by ratio-paired t-test. P-values: *P < 0.05, **P < 0.01, ***P < 0.001, ****P < 0.0001.

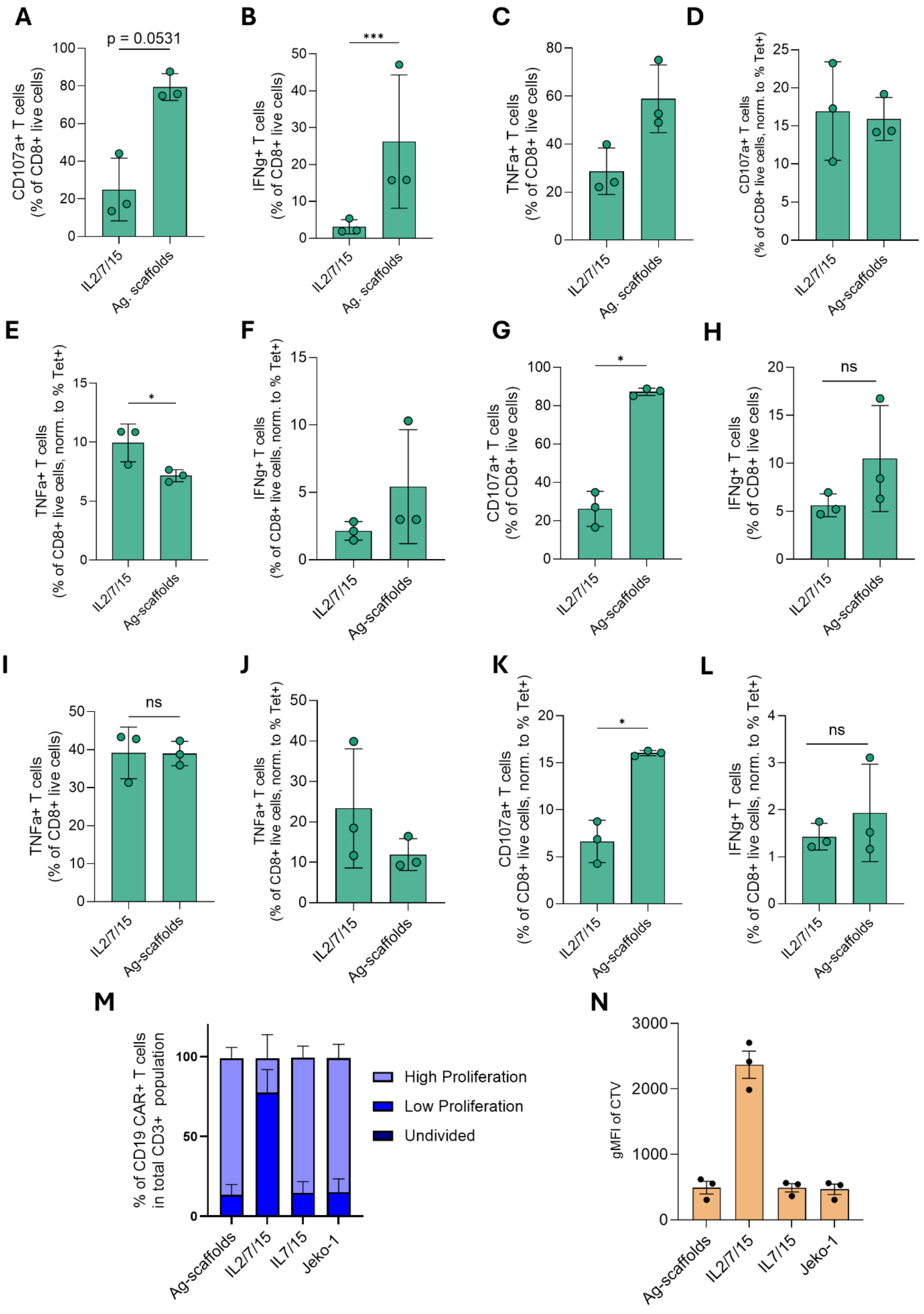
A
p = 0.0531
CD107a+ T cells (% of CD8+ live cells)
IL2/7/15
Ag. scaffolds
B
***
IFNg+ T cells (% of CD8+ live cells)
IL2/7/15
Ag. scaffolds
C
TNFa+ T cells (% of CD8+ live cells)
IL2/7/15
Ag. scaffolds
D
CD107a+ T cells (% of CD8+ live cells, norm. to % Tet+)
IL2/7/15
Ag-scaffolds
E
*
TNFa+ T cells (% of CD8+ live cells, norm. to % Tet+)
IL2/7/15
Ag-scaffolds
F
IFNg+ T cells (% of CD8+ live cells, norm. to % Tet+)
IL2/7/15
Ag-scaffolds
G
*
CD107a+ T cells (% of CD8+ live cells)
IL2/7/15
Ag-scaffolds
H
ns
IFNg+ T cells (% of CD8+ live cells)
IL2/7/15
Ag-scaffolds
I
ns
TNFa+ T cells (% of CD8+ live cells)
IL2/7/15
Ag-scaffolds
J
TNFa+ T cells (% of CD8+ live cells, norm. to % Tet+)
IL2/7/15
Ag-scaffolds
K
*
CD107a+ T cells (% of CD8+ live cells, norm. to % Tet+)
IL2/7/15
Ag-scaffolds
L
ns
IFNg+ T cells (% of CD8+ live cells, norm. to % Tet+)
IL2/7/15
Ag-scaffolds
M
% of CD19 CAR+ T cells in total CD3+ population
High Proliferation
Low Proliferation
Undivided
Ag-scaffolds
IL2/7/15
IL7/15
Jeko-1
N
gMFI of CTV
Ag-scaffolds
IL2/7/15
IL7/15
Jeko-1

**Figure S3. Intracellular cytokine staining in TCR and CAR T cells.** (**A-F**) CRISPR/Cas9 TCR KI T cells expanded using TCR Ag-scaffolds or cytokine cocktail (IL2/7/15). Intracellular staining for CD107a, IFN-γ, TNF-α, 6 hrs after re-stimulation with target cells in 1:1 Effector: Target ratio. **(A-C)** Graph showing the expression % of T cells expressing when re-stimulated with target cells for 6 hrs **(A)** CD107a; **(B)** IFN-γ**;** **(C)** TNF-α; **(D-F)** Graph showing the expression % of T cells expressing when re-stimulated with target cells for 6 hrs normalised to the number of NY-ESO-1-specific 1G4 $TCR^+$ T cells **(D)** CD107a; **(E)** IFN-γ**;** **(F)** TNF-α; **(G-L)** TCR T cells over 12 days post-lentiviral transduction and expansion (MOI-1). Intracellular staining for CD107a, IFN-γ, TNF-α, 6 hrs after re-stimulation with target cells in 1:1 Effector: Target ratio. **(G-I)** Graph showing the expression % of T cells expressing when re-stimulated with target cells for 6hrs **(G)** CD107a; **(H)** IFN-γ**;** **(I)** TNF-α; **(J-L)** Graph showing the expression % of T cells expressing when re-stimulated with target cells for 6 hrs normalised to the number of NY-ESO-1-specific 1G4 $TCR^+$ T cells **(J)** TNF-α; **(H)** CD107a; **(I)** IFN-γ**;** **(M, N)** Proliferative capacity of expanded CRISPR/Cas9 KI CD19 CAR T cells, as measured by loss of Cell Trace Violet in a 6-day co-culture with irradiated Jeko-1 target cells at an Effector: Target ratio of 1:5. **(M)** Proliferative capacity based on cell trace violet expression of CAR T cells on day 6 of restimulation. **(N)** Graph showing geometric MFI of the cell trace violet on day 6. Statistical significance in (**A, B, E, G, H, I, K, L**) determined by ratio-paired t-test: $*P < 0.05$, $**P < 0.01$, $***P < 0.001$, $****P < 0.0001$.

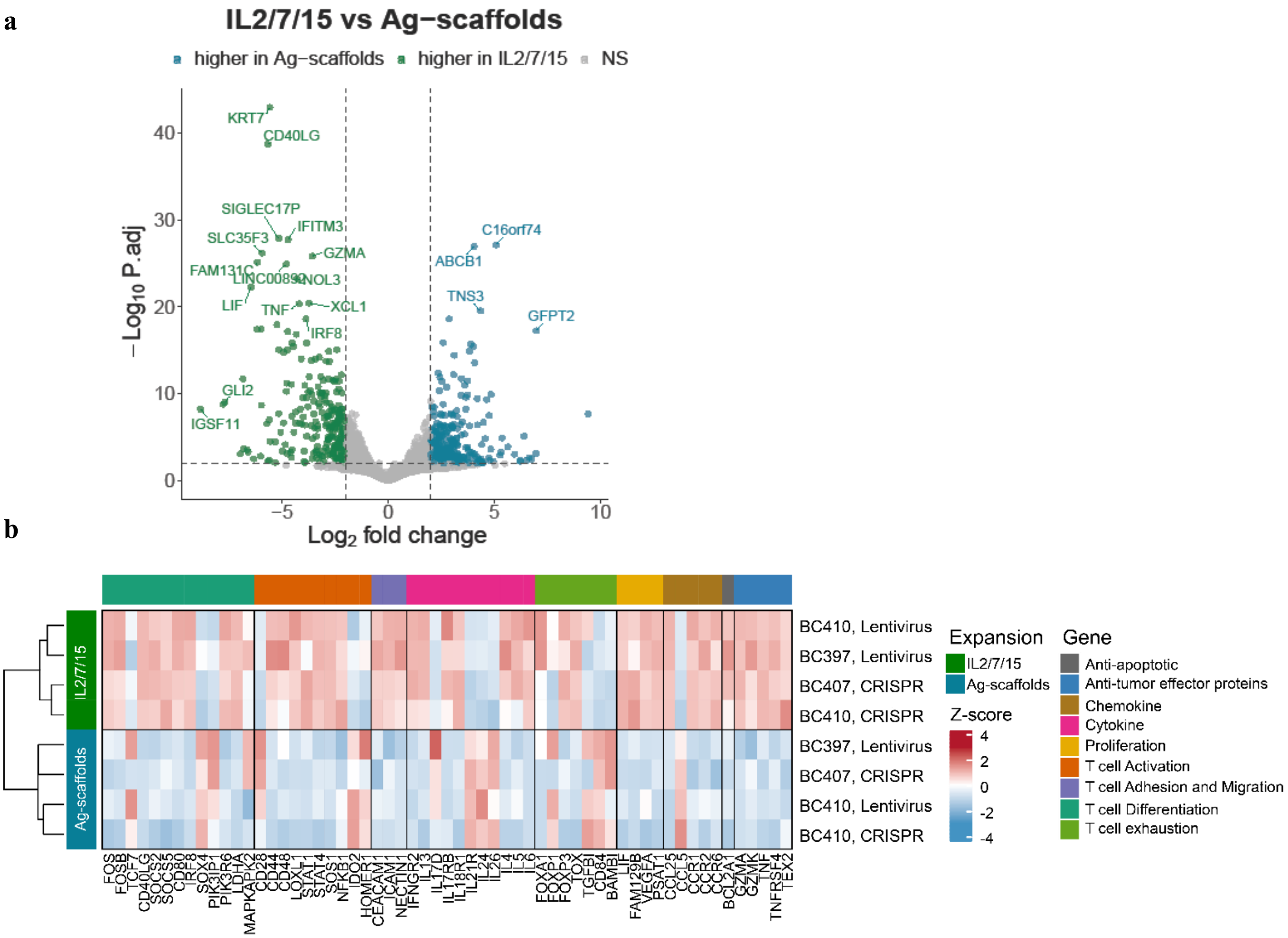


**Figure S4. Bulk RNA sequencing of TCR T cells expanded with Ag-scaffolds or IL2/7/15.** (**A**) Volcano plots showing differentially expressed genes for cells expanded with Ag-scaffold compared to IL2/21 or IL2/7/15. Genes significantly up- or down-regulated in the Ag-scaffold-expanded cluster are highlighted in red and blue, respectively. Dotted lines indicate thresholds on significance (p_threshold = 0.01; log2FC_threshold = 2). (**B**) Differential gene expression in CRISPR/Cas9 or lentiviral TCR T cells expanded with Ag-scaffold or IL2/7/15. Heat map showing expression levels of selected genes.

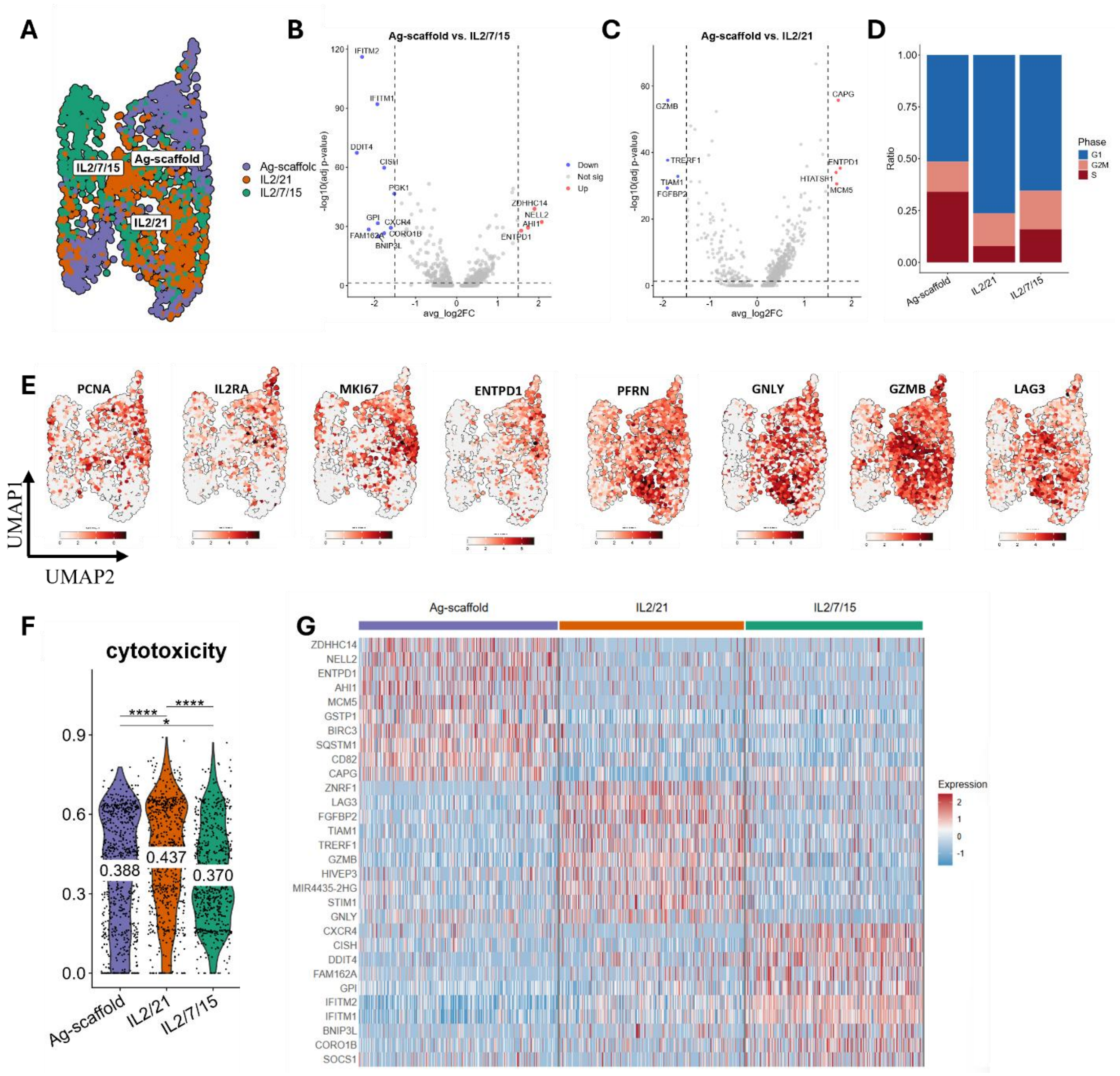


**Figure S5. Single-cell RNA sequencing (scRNAseq) of lentiviral-engineered T cells expanded with either Ag-scaffold, IL2/7, or IL2/7/15.** (**A**) Uniform Manifold Approximation and Projection (UMAP) plot of all lentiviral engineered TCR T cells (total; 2104 cells), with clusters colored and annotated according to cells expanded with either Ag-scaffold (total; 746 cells), IL2/21 (total; 665 cells), or IL2/7/15 (total; 693 cells). (**B**, **C**) Volcano plots showing differentially expressed genes for cells expanded with Ag-scaffold compared to IL2/7/15 (left) or IL2/21 (right). Genes significantly up- or down-regulated in the Ag-scaffold-expanded cluster are highlighted in red and blue, respectively. Dotted lines indicate thresholds on significance (log-fold-change =< 1.5 and p-value < 0.05 (-log10(0.05) = 1.3)). (**D**) Barplot showing the ratio of cells in each cell cycle phase (G1, S, G2/M) across the three expansion conditions. **(E)** UMAP plot of all CRISPR/Cas9 engineered TCR T cells showing the single cell mRNA expression levels of GNLY,

GZMB, LAG3, PCNA, IL2RA, MKI67. (**F**) Violin plot illustrating cytotoxicity gene signature scores and means across the expansion conditions, calculated from RNA expression of selected markers using UCell (Cytotoxicity: GZMA, GZMB, GNLY, PRF1, ITGB1, LAMP1). (**G**) Heatmap of top 10 differentially expressed genes for each group. Statistics are calculated using Wilcoxon signed-rank test. ns = not significant, *P < 0.05, **P < 0.01, ***P < 0.001, ****P < 0.0001.

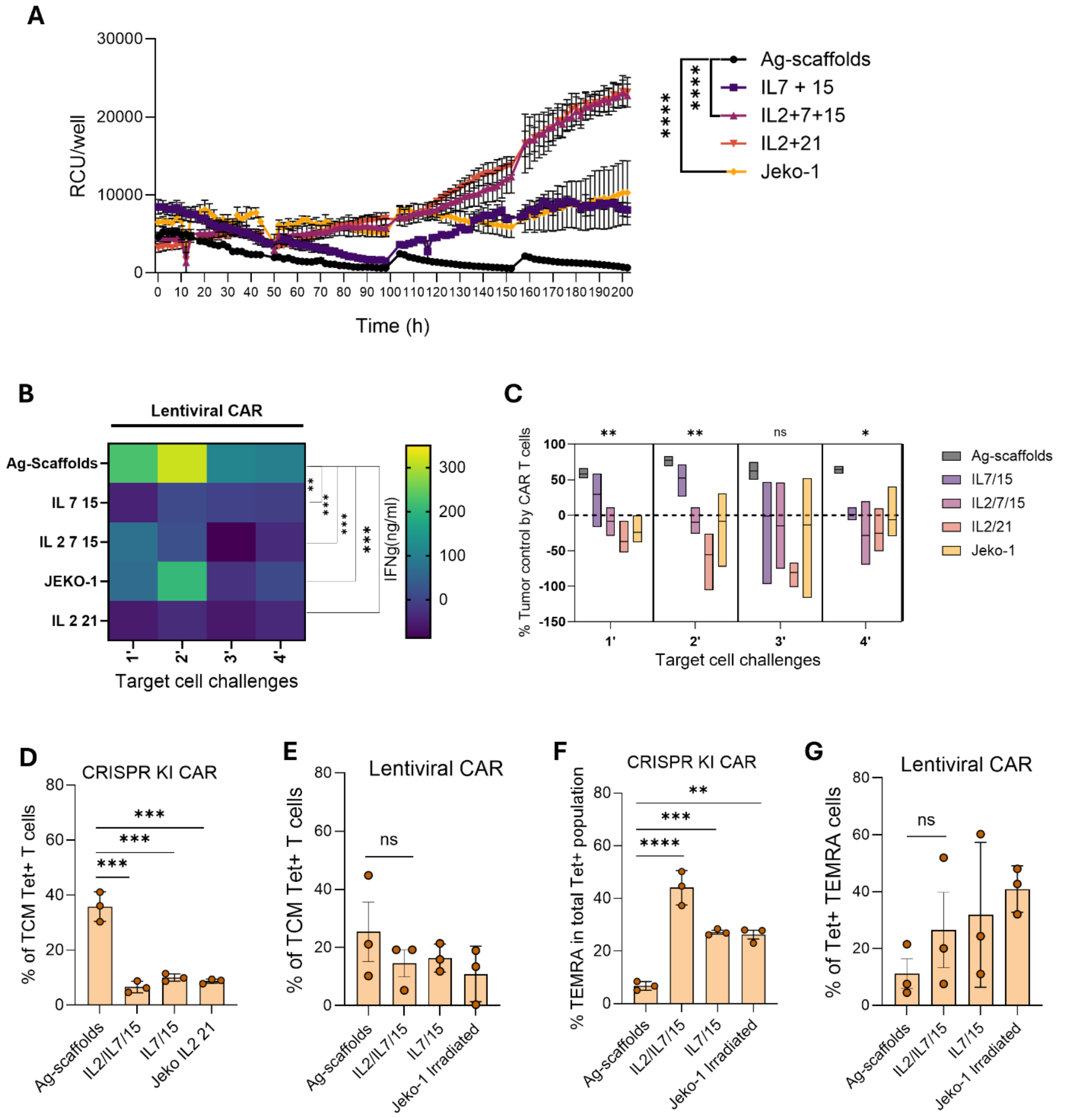


**Figure S6**. **CAR T cells over 12 days post-lentiviral transduction and expansion. (A)** Cytotoxicity of CAR T cells in co-culture with Jeko-1 cells at an Effector: Target ratio of 1:1, tracked over 200 hrs in the Incucyte with rechallenge every 48 hrs, represented by relative confluence units (RCU) per well. **(B)** IFN-γ_expression in the supernatant_(ng/mL) by lentiviral CAR T cells after every 48 hours of target cell challenge. (**C**) Graph showing, percentage tumour control by the CAR T cells over different target cell challenges calculated by %Tumour Control = (RCU at 48 hr – RCU at 0 hr)/(RCU at 0 hr)*100 (n=3 donors from 1 experiment). (**D-E**) Graph shows the percentage of $CD45RA^{-}CCR7^{+}$ central memory CAR T cells in CRISPR KI (**D**) and lentiviral (**E**) CAR T cells respectively assessed after 4 rounds of rechallenge (n=3 donors from 1 experiment). **(F-G)** terminally differentiated $CD45RA^{+}CCR7^{-}$ CRISPR (**F**) or lentiviral (**G**) CAR T cells assessed after 4 rounds of rechallenge (n=3 donors from 1 experiment). Statistical significance in (**A, B, C, D, E, F, G**) was determined by one-way ANOVA with Holm-Šídák's multiple comparisons test, P-values: *P < 0.05, **P < 0.01, ***P < 0.001, ****P < 0.0001.

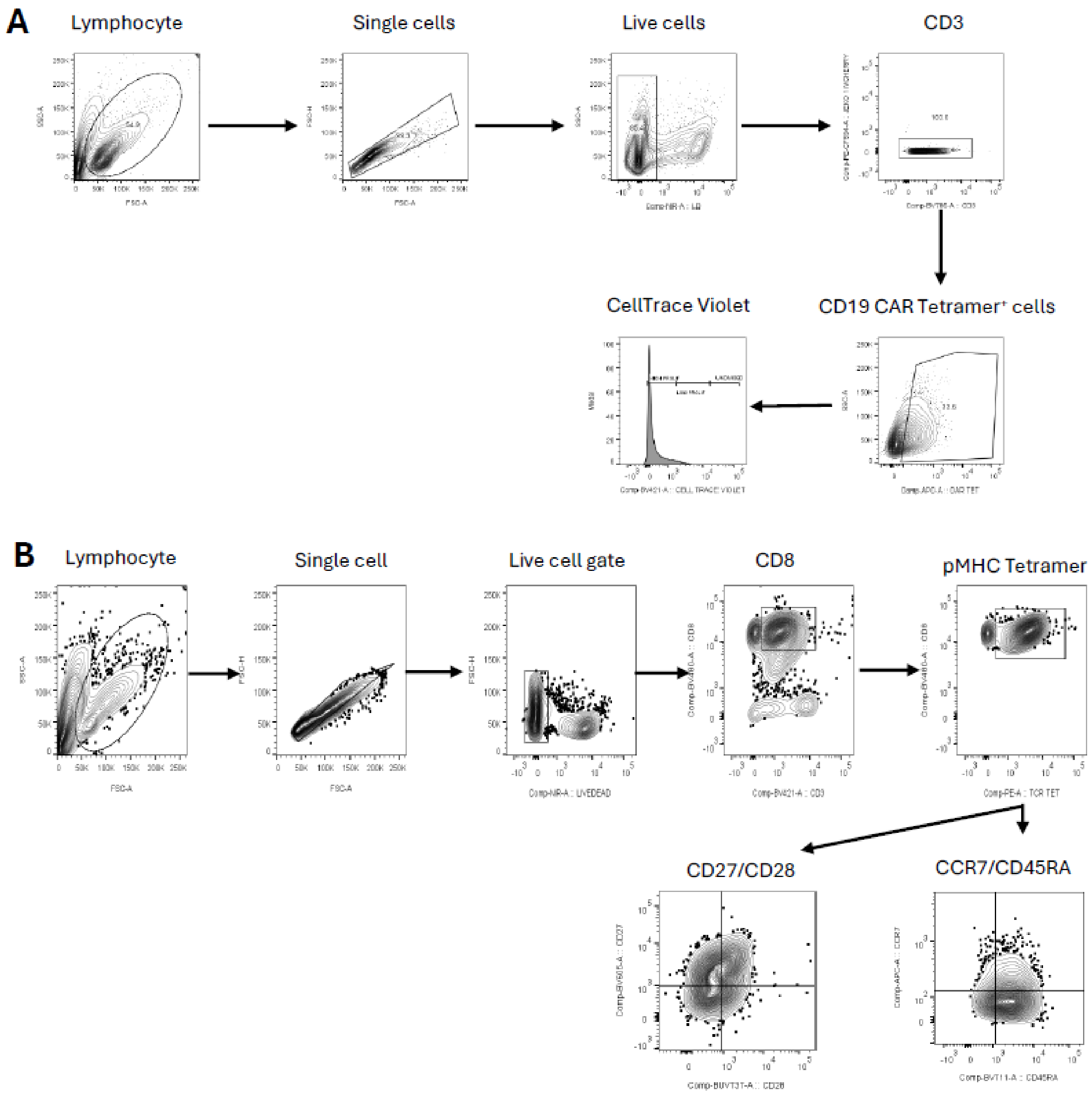


**Figure S7**. **Gating strategy 1.** (**A-B**) Representative gating strategy for (A) restimulation proliferation experiment and (B) T cell phenotype.

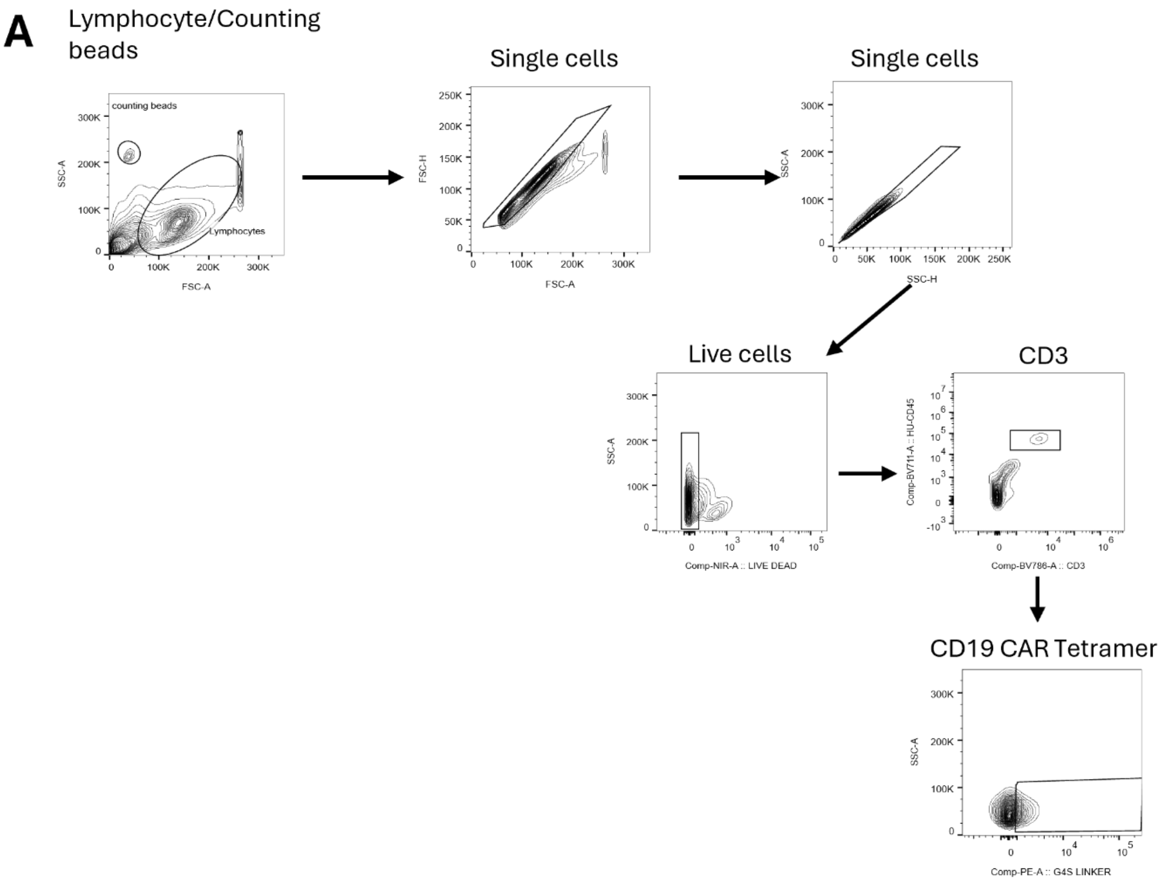


**Figure S8**. **Gating strategy 2.** (**A**) Gating strategy for analysis of mouse spleen and bone marrow.
